# Disentangling spatial organization and splicing of rare intron classes in the human genome

**DOI:** 10.1101/2025.08.11.669784

**Authors:** Saren M. Springer, Katherine Fleck, Kaitlin N. Girardini, Sean M. Riccard, Jelena Erceg, Rahul N. Kanadia

## Abstract

Three-dimensional (3D) genome organization influences transcription and RNA processing, yet how the spatial positioning of genes contributes to pre-mRNA splicing has only recently come into focus. Despite these advances, it remains unclear how introns, particularly rare intron classes, are organized within the 3D genome and whether this organization influences their splicing. Here, we mapped the spatial organization of six intron classes including major, minor, minor-like, hybrid, major-like and non-canonical across four human cell lines (K562, H1, HCT116, and HFFc6) using Hi-C, TSA-seq, and DamID-seq data. This revealed minor intron enrichment in active compartments A and speckle-associated domains (SPADs) and depletion from lamina-associated domains (LADs), whereas hybrid and non-canonical introns showed the opposite trend. Integrating TSA-seq with RNA-seq data suggested that splicing efficiency depends on intron identity rather than nuclear positioning. For example, major-like, minor-like, and non-canonical introns in SPADs were less efficiently spliced than major and minor introns despite their proximity to nuclear speckles. These patterns were consistent across cancer (K562, HCT116) and stem cells (H1) but not fibroblasts (HFFc6). Comparison of minor intron splicing in and out of SPADs across cell lines revealed that, relative to fibroblasts, minor introns outside of SPADs in cancer cells were more efficiently spliced. This suggests that increased efficiency of minor intron splicing in cancer cell lines is not necessarily due to 3D positioning. In all, these findings reveal that intron subclasses show distinct nuclear organization, yet for minor introns, identity rather than position governs splicing efficiency.

## Introduction

The 3D architecture of the genome has emerged as a critical regulatory layer of gene expression, with spatial positioning of genes within the nucleus influencing transcriptional and splicing activity. Since the early distinction between euchromatin (open, active) and heterochromatin (condensed, repressed), the invention of technologies such as Hi-C, TSA-seq, and others have facilitated the systematic mapping of chromatin to nuclear locales and significantly expanded our understanding of the 3D genome (1–7). At a broad scale, chromatin can segregate into two distinct compartments, with compartment A corresponding to open chromatin and transcriptionally active genomic regions, and compartment B corresponding to closed or repressed chromatin (1). These compartments can be further refined into five subcompartments, A1, A2, B1, B2, and B3, associated with distinct histone marks, subnuclear structures, and replication timing (2, 8). At a smaller scale, domains (also known as topologically associating domains; TADs) constitute genomic regions that exhibit high levels of self-interaction, while boundaries with lower levels of chromosomal interactions can insulate adjacent domains (2, 9–11). TADs may be formed through loop extrusion, which brings two linearly distant genomic sites into spatial proximity, termed loop anchors (2, 12, 13). Furthermore, techniques such as TSA-seq, DamID-seq and others, allow for chromatin and genes to be mapped relative to nuclear landmarks such as speckles, nucleoli, and the lamina (3–7). These spatial features are not only descriptive but also actively influence gene regulation, reinforcing the idea that co-localized genes may share regulatory mechanisms (2–7, 14–32). Here, we investigate how intron classes are organized within the 3D genome and whether this positioning informs their splicing.

Introns originated in the last eukaryotic common ancestor and have persisted for 1.8 billion years, expanding markedly in metazoans, which we and others have hypothesized to play a significant role in regulating multicellular complexity (33–39). Their presence and removal from pre-messenger RNAs necessitated the co-evolution of the spliceosome, whose snRNA-guided base-pairing and two transesterification reactions excise introns and join exons (40–42). Despite being non-coding, introns harbor essential splicing motifs, including the 5′ and 3′ splice sites, branch point sequence, and polypyrimidine tract (43, 44). Based on these motifs and their relative abundances, introns have traditionally been classified as major or minor. However, the observation of divergent consensus sequences led us to reclassify introns along a continuum spanning major, major-like, hybrid, minor-like, minor, and non-canonical classes, raising the question of how these rare intron classes are spliced (45–50). While the abundance of major introns and high levels of major snRNAs inherently facilitates their splicing, recent work has shown that major intron splicing is not purely stochastic as recruitment to nuclear bodies, such as speckles, actively promotes their splicing (31, 32, 43, 51–63). By contrast, minor introns and their corresponding snRNAs are lowly abundant, leading to the idea that minor introns are a rate-limiting step in minor intron-containing gene (MIG) expression and thus prone to intron retention (49, 64–69). However, we and others have shown that minor intron retention is a regulated outcome, not merely a sign of inefficiency (68, 70–73). Thus, we hypothesize that 3D genome architecture may organize minor introns and other rare intron classes into spatially privileged regions to ensure proper splicing.

To test this hypothesis, we leveraged Hi-C, TSA-seq, DamID-seq, and RNA-seq datasets from K562 cells, a human chronic myelogenous leukemia cell line commonly used to study 3D genome organization (2, 3, 5, 8, 74–80). We mapped six intron classes, major, major-like, hybrid, minor-like, minor, and non-canonical, across nuclear compartments, subcompartments, nuclear bodies, and 3D genomic features. Despite their low abundance, rare intron classes, including minor, minor-like, hybrid, and non-canonical, showed distinct spatial biases within the 3D genome. Minor introns were highly enriched in both compartment A and SPADs, which are linked to active transcription and splicing, respectively (31, 32). Integration with transcriptome data revealed that minor introns are frequently found in expressed genes. Consistently, few minor introns overlapped with repressive regions, as such statistical analysis was not possible in some cases. However, sufficient intron numbers in SPADs provided an opportunity to assess splicing efficiency across intron classes. Within SPADs, major and minor introns were efficiently spliced, while other rare intron classes showed reduced splicing efficiency. This trend was consistent across K562 and three additional cell lines encompassing human fibroblast cells (HFFc6), embryonic stem cells (H1), and colorectal cancer cells (HCT116). Comparing splicing across four cell lines, we found that while major intron splicing in SPADs varied, minor introns consistently exhibited efficient splicing. Using non-cancerous human fibroblasts as a reference, we found the splicing of matched minor introns from cancer cell lines was more efficient when outside of SPADs. Functional enrichment of MIGs with enhanced splicing in both cancer cell lines revealed an overrepresentation of cell cycle regulators. Thus, in cancer cells, proximity to SPADs is not required for efficient splicing of minor introns, which aligns with our previous finding that minor spliceosome snRNAs are upregulated in cancer cells to meet the high demands for MIG-controlled functions (81). Together, our results suggest that minor intron splicing is buffered against spatial constraints, implying a form of regulatory resilience and raising new questions about how rare introns are accommodated within the 3D genome.

## Results

### Rare introns are scattered across the linear human genome

Across the human genome, exons comprise only 3.83% of gene-body sequence, whereas 96.17% is occupied by introns **(Fig. 1A, Table S1)**. Amongst these, 93.99% are major introns (345,317), while the remaining 6.01% consist of rare intron classes. In order of abundance, these are: major-like (29,337; 5.03%), non-canonical (6070; 0.74%), minor (850; 0.11%), minor-like (458; 0.06%), and hybrid (373; 0.07%) introns **(Fig. 1A and B, Fig. S1A, Table S1)**. Given this disparity in abundance, we investigated whether the distribution of rare introns reflected their frequency or showed enrichment within specific clusters. We first examined inter-intron distance i.e., the distance between two consecutive introns of the same class. We reasoned that if rare introns are found near one another, they may share common regulatory mechanisms. Furthermore, genes that contain rare introns usually harbor only one, with all remaining introns being major **(Fig. S1B and C).** Consequently, rare introns of the same class are generally located in different genes, predicting greater inter-intron distances than those observed for abundant major introns. Indeed, we found that as intron abundance decreased, inter-intron distance increased **(Fig. 1C and D)**. Notably, the median inter-intron distance for major introns was 141 bp. Given the abundance of major introns, it is not surprising that the inter-intron distance is approximately the same as the median human exon length of 120 bp **(Fig. 1C)** (82). The next most abundant intron class was major-like which showed a median inter-intron distance of 27 kb, followed by 105 kb for non-canonical introns. This distance for minor introns was 1.8e06 bp and for minor-like was 4.0e06 bp. Finally, hybrid introns, which are the rarest of the subclasses, had an inter-intron distance of 6.1e06 bp **(Fig. 1C)**. The progressive increase in the inter-intron distance with the rarity of intron class was investigated by correlation analysis, which revealed a nearly linear relationship between inter-intron distance and intron abundance **(Fig. 1D; R^2^ = 0.98)**.

**Figure 1.**
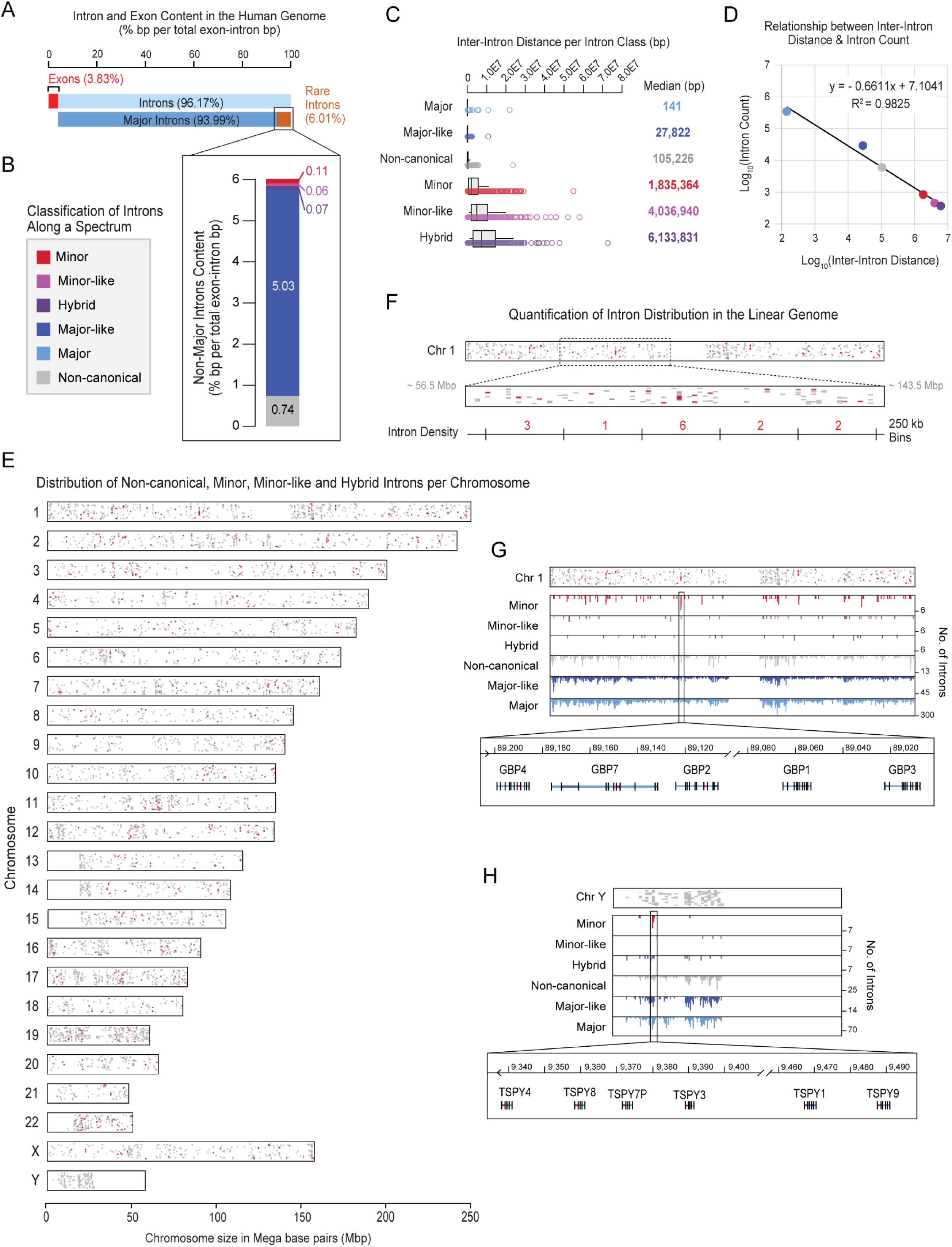
Distribution of intron classes across the linear genome. A. Bar graph showing the percentage of the coding and non-coding genome occupied by exons, major introns, and rare introns. B. Bar graph showing the percentage of the coding genome occupied by rare intron classes, including minor, minor-like, hybrid, major-like, and non-canonical introns, as defined in Olthof-Schwoerer et al. (49) C. Boxplot showing the distribution of inter-intron distance (in base pairs, bp) for each intron class. Median distance per class is listed to the right of each boxplot. D. Scatter plot establishing the relationship between inter-intron distance and intron number per class. Line of best fit and R^2^ value are shown in the graph area. E. Chromosomal map depicting the genome-wide distribution of non-canonical, minor, minor-like, and hybrid introns. Introns are color coded in accordance with the spectrum listed in panel B. F. Zoom-in of Chromosome 1 from panel E. Visual summary of depicting quantification of intron density in fixed genomic windows. Approximate genomic location is shown in mega bp (Mbp). G. Intron density per 250 kb bin along chromosome 1. Inset highlights a gene cluster located within a single 250 kb bin composed of multiple MIGs from the GBP gene family. H. Intron density per 250 kb bin along the Y chromosome. Inset highlights a gene cluster located within a single 250 kb bin composed of multiple MIGs from the TSPY gene family.

While the rarity of an intron is related to the inter-intron distance, we explored the possibility that there might be clusters of introns within specific chromosomes that get diluted in this overall analysis. Therefore, we visualized rare intron density per chromosome by plotting minor, minor-like, hybrid, and non-canonical introns within a chromoMap **(Fig. 1E)**. In keeping with our findings of inter-intron distance and abundance, we generally found minor, minor-like, hybrid, and non-canonical introns to be distributed throughout the genome **(Fig. 1E, Fig. S1D-F, Table S2)**. However, there were regions in chromosomes 1, 11, 17, 19, 22, and Y that appeared to have a higher density of rare intron classes. To investigate whether this was a statistically significant phenomenon, we binned the human genome into 250 kb segments **(Fig. 1F, Fig. S2)**, which revealed that rare introns are broadly distributed across all chromosomes without enrichment in any loci—except for two clusters of minor introns: the GBP gene family on chromosome 1 **(Fig. 1G)** and the TSPY gene family on the Y-chromosome **(Fig.1H, Dataset S1)**. Dispersal of rare intron classes, in particular minor introns, across the linear genome raises a key question: how does the minor spliceosome, given its low abundance, locate these rare introns? Therefore, we explored whether minor and other rare intron classes exhibit distinct spatial localization that could facilitate their splicing.

### Intron classes show distinct spatial biases, notably the enrichment of minor introns in compartment A

Using Hi-C data from K562 cells, we examined the spatial distribution of intron subtypes across various 3D features of the nucleus, such as compartments A, B, and unassigned regions (“neither”) **(Fig. 2A, Datasets S2-S6)**. We leveraged the abundance of major introns in the human genome as a background distribution of introns within compartments, which served as a reference for comparing the distribution of rare introns. For example, 62% of major introns were found in compartment A, while 64% of major-like introns **(Fig. 2B, Fig. S3A; p = 4.2e-16),** and 71% of minor introns were significantly enriched in compartment A **(Fig. 2B and C; p = 4.7e-08)**. In contrast, minor-like introns showed no significant difference from major introns **(Fig. 2B, Fig. S3A)**. For classes whose distributions shifted relative to major introns, we observed reciprocal patterns, where gains in one compartment mirrored losses in another. For example, minor introns, which showed significant enrichment in compartment A, showed a reciprocal reduction in compartment B relative to major introns **(Fig. 2C; p = 3.2e-10)**. We observed that 51% of hybrid **(Fig. 2B, Fig. S3A; p = 1.8e-05)** and 50% of non-canonical **(Fig. 2B, Fig. S3A; p = 2.2e-16)** introns were significantly underrepresented in compartment A. Instead of the expected increase of these intron classes in compartment B, we found that 17% of hybrid **(Fig. 2B, Fig. S3A; p = 4.2e-16)** and 22% of non-canonical **(Fig. 2B, Fig. S3A; p = 2.2e-16)** introns were enriched in the “neither” category. Although multiple intron classes exhibited spatial bias, minor introns showed the strongest enrichment in compartment A, suggesting a non-random spatial organization despite their rarity.

**Figure 2.**
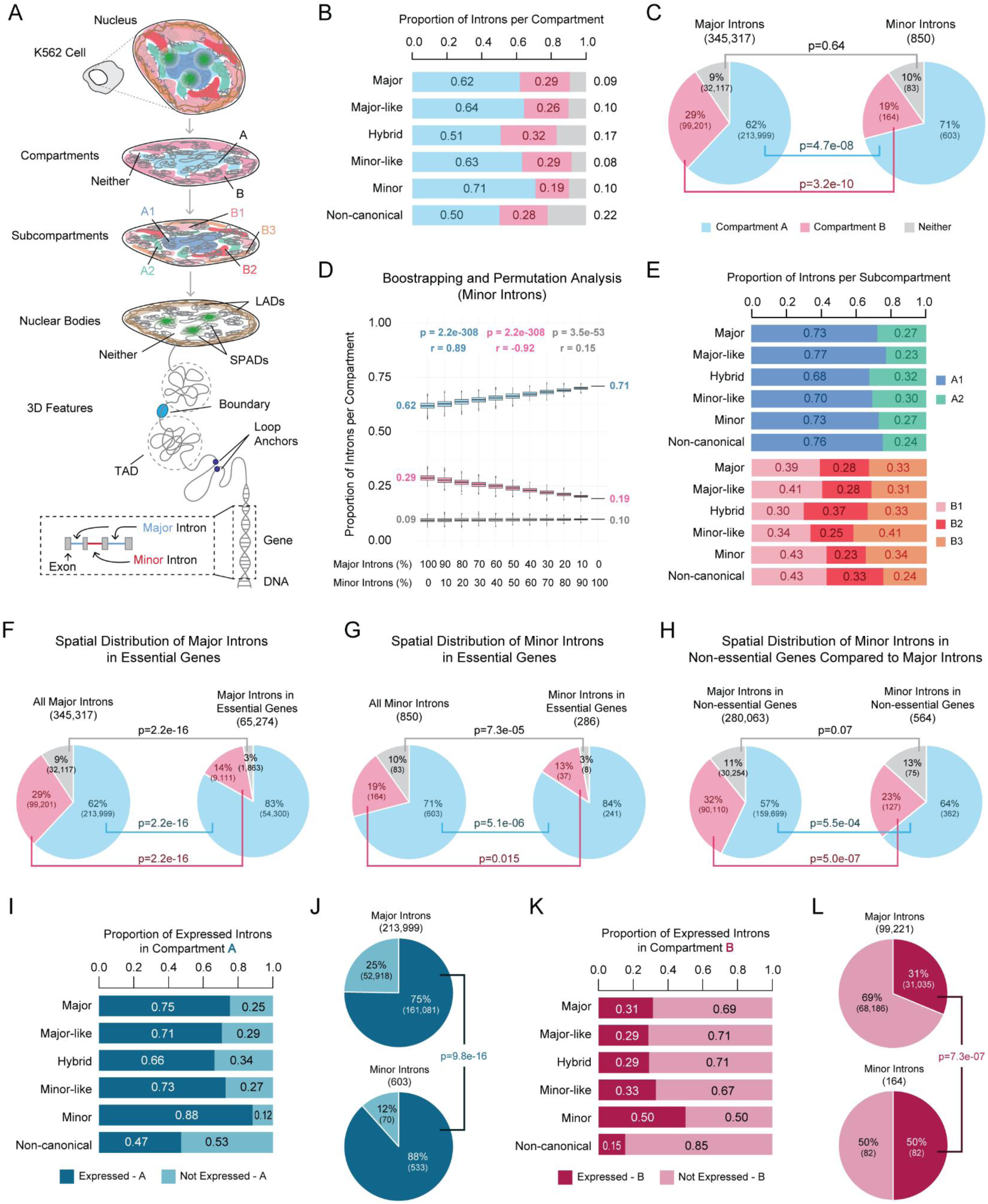
Introns exhibit class-specific biases in 3D genome organization. A. Schematic showing the levels of genome organization interrogated in this study. Annotations were derived from K562 cells and include compartments (A and B), subcompartments (A1, A2, B1, B2, and B3), domains associated with nuclear bodies (SPADs and LADs), and 3D features (TADs, boundaries, and loop anchors). B. Proportional distribution of each intron class across compartments A, B, and unassigned regions (“neither”). C. Fisher’s exact test comparing the compartment distribution of major versus minor introns. D. Bootstrapping and permutation analysis depicting the proportion of introns per compartment across 1000 bootstrapped intron sets that vary the percentage of major introns versus minor introns (see also Fig. S4). E. Proportional distribution of introns localized to compartments A and B across their respective subcompartments (A1, A2, B1, B2, and B3), shown for each intron class. (F-H) Fisher’s exact test comparing the compartment distribution of (F) all major introns versus major introns in essential genes, (G) all minor introns versus minor introns in essential genes, and (H) major introns versus minor introns in non-essential genes. I. Proportion of introns per class that are in compartment A and fall within expressed genes. J. Fisher’s exact test comparing the distribution of major versus minor introns in compartment A among expressed genes. K. Proportion of introns per class that are in compartment B and fall within expressed genes. L. Fisher’s exact test comparing the distribution of major versus minor introns in compartment B among expressed genes.

While the enrichment of minor introns in compartment A suggests biological relevance, a key confounder is that intron class inherently selects for a specific subset of genes, potentially biasing spatial localization. In other words, spatial enrichment of rare introns may reflect localization of the genes they reside in, rather than properties inherent to the intron itself. To address this, we performed bootstrapping and permutation analyses, generating 1,000 random sets of 850 major introns, to match the number of minor introns, and evaluated their enrichment in compartments A and B. The median enrichment in compartment A for 1,000 lists of 850 major introns remained at 62%, identical to the full major intron set **(Fig. 2D)**, suggesting the observed minor intron enrichment of 71% is not an artifact of gene selection. To test whether the significant enrichment we observed for minor introns in compartment A was tethered to intron identity, we next performed pollution analysis. Here, we took the 1,000 major intron lists and substituted a portion of major introns with minor introns (in increments of 10%…100%). This analysis revealed a progressive increase in intron enrichment from 10 to 100% minor intron composition, supporting the idea that minor introns are highly enriched in compartment A **(Fig. 2D; r = 0.89, p = 2.2e-308)**.

Performing bootstrapping and pollution analyses on other intron classes in compartment A, B, or the “neither” category revealed the observed enrichment was indeed tethered to their identity and not their number and/or gene subtype **(Fig. S4)**. These results confirm that the 3D positioning of rare introns is not a byproduct of gene identity or intron abundance but instead reflects class-specific compartmental organization.

We further refined compartmental resolution by subdividing compartment A into A1 and A2, and compartment B into B1, B2, and B3 **(Fig. 2A)**. This allowed us to determine whether enrichment in compartments A or B stemmed from specific subcompartments. We found that, of the introns in compartment A, 77% of major-like **(Fig. 2E, Fig. S3B; p = 2.2e-16)** and 76% of non-canonical **(Fig. 2E, Fig. S3B; p = 9.5e-05)** introns were enriched in A1, which was significantly higher than the 73% of major introns in A1 **(Fig. 2E, Fig. S3B)**. Other rare intron classes showed no differences in enrichment between A1 and A2 **(Fig. S3B)**. Across B1-3, all intron classes—except minor— differed significantly from the major intron distribution (B1: 39%, B2: 28%, B3: 33%) **(Fig. S3C)**. Major-like introns were enriched in the more facultative heterochromatin B1 subcompartment **(Fig. S3C; 41%, p = 2.2e-16)** and depleted from the peripheral B3 subcompartment **(Fig. S3C; 31%, p = 2.2e-16)**. Minor-like introns showed significant increases in B3 **(Fig. S3C; 41%, p = 0.031)** but not in B1 **(Fig. S3C; 34%, p = 0.24)** and B2 **(Fig. S3C; 25%, p = 0.38)**. Hybrid introns were depleted in B1 **(Fig. S3C; 30%, p = 0.039)** and enriched in B2 **(Fig. S3C; 37%, p = 0.042)**. Non-canonical introns were enriched in B1 **(Fig. S3C; 43%, p = 1.2e-03)** and B2 **(Fig. S3C; 33%, p = 1.5e-04)** but significantly depleted from B3 **(Fig. S3C; 24%, p = 3.6e-13)**. To test whether the observed biases were tethered to intron identity, we again performed bootstrapping and pollution analyses for subcompartment distributions. We found that the subcompartment distribution of rare intron classes is not a byproduct of gene identity or abundance **(Fig. S5)**. Finally, we explored the distribution of introns at a finer resolution, such as TADs, loop anchors, and boundaries. Assessing intron distribution in combinations of 3D genomic features revealed that only non-canonical introns were significantly enriched in loop anchors and boundaries, underscoring their unique spatial organization **(Fig. S6, Table S3)**.

### Minor intron enrichment in compartment A is partially independent of their enrichment in essential genes

We have previously shown that minor introns are enriched in essential genes (37), which are frequently found at sites of active gene expression in the nuclear interior, such as near nuclear speckles (3–5, 83). Thus, we hypothesized that the essential nature of MIGs could contribute to their enrichment in compartment A. Given our recent updates to minor intron annotations and the addition of new intron classes, we revisited our baseline enrichment of each intron class in essential genes (49). For this, we leveraged our previously established essentialome annotations, comprising genes required for the survival of 341 cancer cell lines **(Dataset S7)** (37, 84). The essentialome includes two categories: the total essentialome (genes essential in at least one cell line) and the core essentialome (genes essential in all cell lines). Minor intron-containing genes were significantly enriched in both the total and core essentialome **(Fig. S7A-C)**. Similarly, major-like intron-containing genes and minor-like intron-containing genes were significantly enriched in both the total and core essentialome **(Fig. S7D-F, S7J-L)**. Genes containing hybrid introns showed enrichment in the total, but not core, essentialome **(Fig. S7G-I)**. Finally, non-canonical intron-containing genes showed significant depletion from essential genes **(Fig. S7M-O)**.

To assess whether essentiality contributes to compartment localization for rare intron classes, we next intersected introns in essential genes with compartment annotations **(Fig. 2F-H, Fig. S8A)**. We found that 83% of major introns in essential genes were enriched in compartment A, which was statistically higher than the overall major intron enrichment in compartment A **(Fig. 2F; p = 2.2e-16)**. Similarly, 84% of minor introns in essential genes were enriched in compartment A, which was statistically higher than the overall enrichment of minor introns in compartment A **(Fig. 2G; p = 5.1e-06)**. These findings are consistent with prior reports and extended to the other rare intron classes, as every intron class had statistically elevated enrichment in compartment A when comparing introns in essential genes to all introns per class **(Fig. S8B)** (83). Given the enrichment of essential MIGs in compartment A, we next wondered if non-essential MIGs were depleted from compartment A. Intersection of non-essential genes containing each intron class with compartment annotations revealed, relative to their overall proportions, a smaller proportion of non-essential genes overlap with compartment A for every intron class **(Fig. S8C)**. However, when the distribution of major introns in non-essential genes is used as a baseline, we found that minor introns in non-essential genes were more highly enriched in compartment A **(Fig. 2H; p = 5.5e-04)**. These findings indicate that minor intron enrichment in compartment A is not solely driven by their presence in essential genes but instead reflects a class-specific spatial bias.

### Minor introns in both compartments A and B are more frequently found in expressed genes

Given that compartment A is generally associated with increased transcriptional activity, we hypothesized that the enrichment of minor introns in compartment A would correlate with MIG expression (1, 25). To test this hypothesis, we leveraged RNA-seq data **(Dataset S8 and S9)**, which revealed that 88% of minor introns in compartment A were in expressed MIGs. This enrichment was statistically higher when compared to the 75% of major introns expressed in compartment A **(Fig. 2I and J; p = 9.8e-16)**. In contrast, 69% of the major introns in compartment B were in genes that were not expressed **(Fig. 2K)**. The same was not true for minor introns in compartment B, as 50% were found in expressed genes **(Fig. 2K and L)**. This enrichment was significantly higher than major introns (31%, p = 7.3e-07), suggesting that MIGs can escape the generally repressed environment of compartment B. Other rare intron classes did not show similar enrichment in expressed genes in compartment A **(Fig. 2I, Fig. S9)**. However, major-like and non-canonical introns in compartment B were significantly less likely to reside in expressed genes compared to major introns **(Fig. 2K, Fig. S9)**. The enrichment of minor introns in expressed genes was also reflected across the subcompartments **(Fig. S10, Dataset S10)**. We also noted significant depletion of non-canonical introns in expressed genes for more refined features, especially boundaries and loop anchors **(Fig. S11)**.

### Minor introns in both SPADs and LADs are more frequently found in expressed genes

Previous studies have reported that compartment A and chromatin around nuclear speckles share a high degree of overlap within the 3D genome (4), a finding we confirmed in our analysis **(Fig. S12A-C)**. Nuclear speckles are membraneless nuclear bodies enriched in spliceosomal proteins, and are defined by the presence of two scaffold proteins, SON and SRRM2 (85), among many other proteins and non-coding RNAs (57, 60, 86–89). Once considered storage sites for splicing factors, nuclear speckles are now recognized as regulatory hubs that influence both transcription and splicing (4, 32, 59, 62, 63, 90, 91). Given the observed enrichment of minor introns and expressed MIGs in compartment A, we next asked whether minor introns were also enriched near nuclear speckles. For this, we leveraged previously generated TSA-seq data using SON, which labeled SPADs in K562 cells (3). This dataset revealed that 25% of minor introns were found in SPADs, a significantly higher percentage than the 21% of major introns in SPADs **(Fig. 3A and B; p = 0.018)**. We observed a reciprocal depletion of minor introns in more peripheral LADs defined using DamID-seq data (5), where only 14% of minor introns were found in LADs compared to 19% of major introns **(Fig. 3A and B; p = 7.7e-05)**. Similarly, only 17% of major-like introns were found in LADs, while 27% were found in SPADs **(Fig. S12D; p = 2.2e-16)**. Minor-like introns did not differ significantly from the major intron distribution within SPADs and LADs **(Fig. S12D)**. However, hybrid and non-canonical introns were significantly underrepresented in SPADs at 9% **(Fig. S12D; p = 5.3e-10)** and 17% **(Fig. 3B; p = 1.8e-15)**, respectively. Correspondingly, 31% of hybrid and 25% of non-canonical introns were found in LADs, a significant overrepresentation compared to major introns **(Fig. 3B, Fig. S12D).**

**Figure 3.**
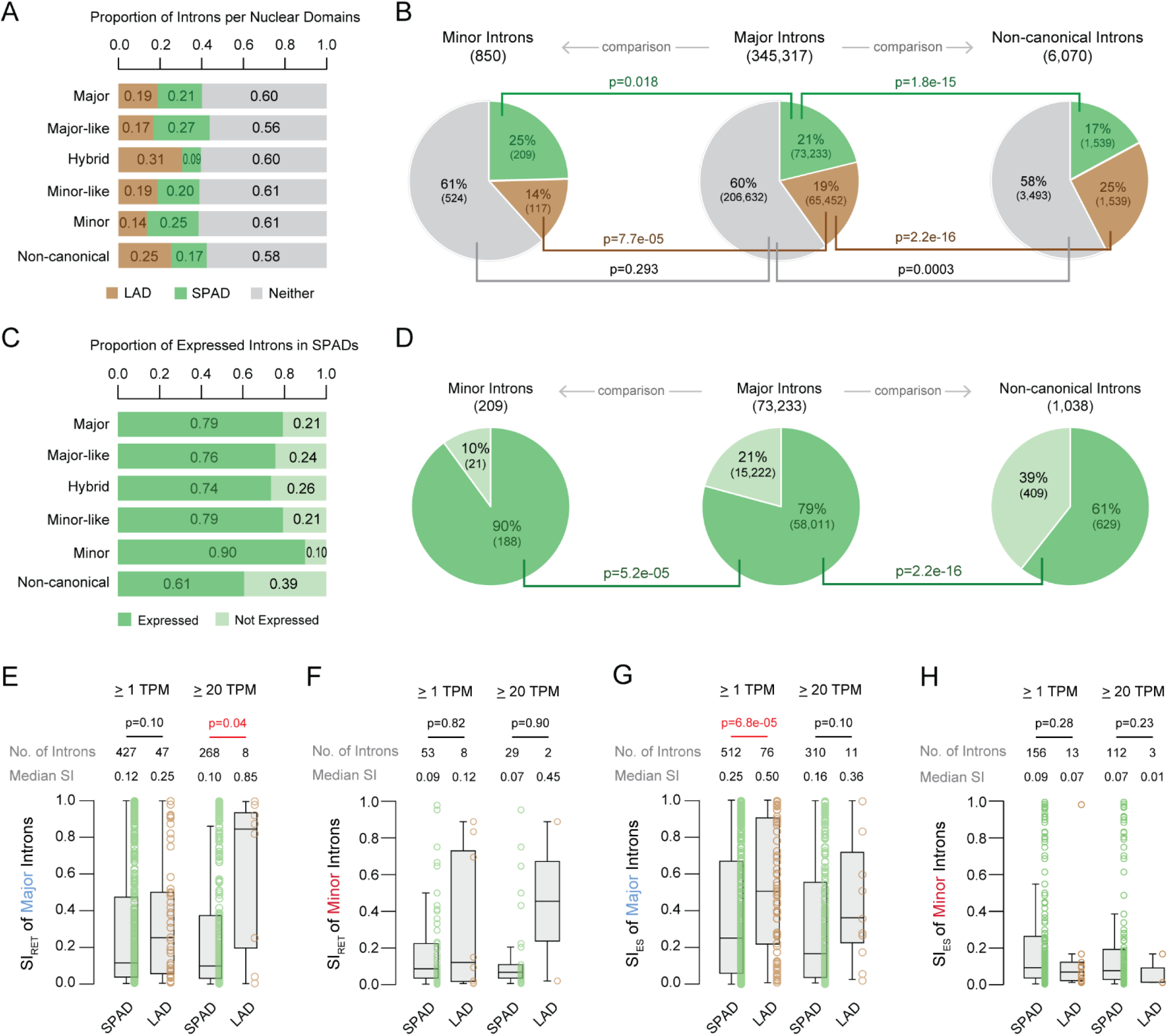
Minor introns are enriched in SPADs, but this enrichment does not correlate with an increase in splicing efficiency relative to LADs. A. Proportional distribution of each intron class across domains associated with nuclear bodies including SPADs, LADs, and unassigned regions (“neither”). B. Fisher’s exact test comparing the nuclear domain distribution of major versus minor introns and major versus non-canonical introns. C. Proportion of expressed introns per class in SPADs. D. Fisher’s exact test comparing the distribution of major versus minor introns and major versus non-canonical introns in SPADs among expressed genes. E-F. Boxplots of major (E) and minor (F) intron retention for genes filtered at ≥ 1 or 20 TPM. G-H. Boxplots of major (G) and minor (H) exon skipping for genes filtered at ≥ 1 or 20 TPM.

Given the regulatory role of nuclear speckles, we hypothesized that rare introns in SPADs may be more likely to be in expressed genes (32, 88). Indeed, 90% of minor introns in SPADs were also in expressed genes, which was significantly higher than major introns at 79% **(Fig. 3C and D; p = 5.2e-05)**. Consistent with our previous observations that more MIGs were expressed in both A and B compartments, we also observed 30% of minor introns in LADs were in expressed genes compared to 18% of major introns (**Fig. S13A and B; p = 0.002)**. The proportion of minor-like and hybrid introns in expressed genes in SPADs or LADs was not significantly different from that of major introns **(Fig. S13C and D)**. However, of the non-canonical introns in SPADs, only 61% were in expressed genes, which is significantly less than major introns **(Fig. 3D; p = 2.2e-16)**. Consistently, non-canonical introns in LADs were also less likely to be in expressed genes at 7%, relative to 18% of major introns **(Fig. S13B; p = 2.2e-16)**. Together, these results suggest that the spatial organization of rare introns within SPADs and LADs is a non-random event and indicates a potential mechanism of gene regulation.

### Minor intron splicing is efficient in SPADs

Next, we explored whether enrichment of minor introns in SPADs led to more efficient splicing. For this, we analyzed minor intron retention in SPADs with modification of our previous bioinformatics strategy to assess intron retention (68, 81, 92–95). We have previously required reads that map to the exon/intron boundary and 90% read coverage across the length of the intron. This conservative mode of assessing intron retention works when we have deep coverage with ribodepleted total RNA-seq. However, in the current study, the K562 transcriptome data is not ribodepleted total RNA-seq, but polyA-enriched mRNA-seq (79). Therefore, we removed the 90% coverage requirement and considered an intron to be retained if we found reads that map to the exon/intron boundary without any evidence of cryptic splicing. In other words, reads that map to an exon/intron boundary are considered as a representation of intron retention, only if there is no evidence of an alternative splicing event that uses cryptic 5’ and 3’ splice sites **(Fig. S14 and S15, Table S4)**.

When applying a minimum expression threshold of 1 transcript per million (TPM), major introns exhibited a non-significant (p = 0.1) trend toward increased intron retention in LADs relative to SPADs **(Fig. 3E)**. This difference reached statistical significance when expression levels were filtered to ≥ 20 TPM, indicating that major intron splicing is less efficient in LADs for more highly expressed genes. This finding is consistent with previous reports linking gene expression levels and spatial localization to splicing outcomes (32, 88). Thus, confirming that our modification to our intron retention bioinformatics pipeline was sufficient to assess splicing efficiency. Applying this strategy to minor introns revealed no significant differences in minor intron retention across SPADs and LADs, regardless of expression threshold **(Fig. 3F)**. This result, while surprising, is not unexpected given that there are only 8 and 2 minor introns in LADs based on ≥ 1 TPM thresholding and ≥ 20 TPM thresholding, respectively. Thus, while minor intron retention was not significantly different between SPADs and LADs, this result should be interpreted with caution due to the limited number of introns analyzed.

Since most minor introns are flanked by major introns, the expression of MIGs requires the coordinated action of both the major and minor spliceosomes (96–98). In fact, we have previously reported that inhibition of the minor spliceosome, besides resulting in intron retention, can in some cases also result in exon skipping, executed by the major spliceosome (68, 95). Thus, we coupled our investigation of intron retention with exon skipping. For exon skipping, major introns in genes expressed at ≥ 1 TPM showed significantly higher skipping in LADs compared to SPADs, though this significance was lost at the ≥ 20 TPM threshold **(Fig. 3G)**. Like intron retention, minor introns showed no significant differences in exon skipping across SPADs and LADs, regardless of expression threshold **(Fig. 3H)**. Notably, among the 13 minor introns in MIGs with a ≥ 1 TPM expression in LADs, exon skipping was generally lower, with one outlier driving variation **(Fig. 3H)**. These results indicate that minor intron splicing in SPADs is efficient, consistent with their enrichment in SPADs and depletion from LADs. Furthermore, minor intron splicing appears to be less affected by lamina proximity than major intron splicing; however, the small number of lamina-proximal MIGs limited further statistical testing.

### Rare classes of introns within SPADs show differential intron retention and exon skipping

Although rare introns were depleted from LADs, their presence in SPADs provided a unique opportunity to compare splicing efficiency across intron classes within a shared spatial context. In K562 cells, comparison of splicing index (SI_RET_) values between intron classes in SPADs revealed that major-like intron showed higher levels of intron retention than major and minor introns **(Fig. 4A)**. Minor-like introns also had a significantly higher median SI_RET_ value than minor introns but did not differ from major introns. Non-canonical introns exhibited more intron retention than major, major-like, and minor introns, but were not significantly different from hybrid or minor-like introns **(Fig. 4A, Dataset S11)**. Only six hybrid introns in SPADs had sufficient data for splicing analysis, although they trended toward inefficient splicing. Of note, minor introns trended toward more efficient splicing than major introns and though this finding was not statistically significant **(Fig. 4A, Dataset S11)**, it is consistent with significantly lower overall intron retention levels for minor introns compared to major introns **(Table S4; p = 0.044).** Similar trends were observed for exon skipping in K562 cells **(Fig. 4B, Dataset S12)**. Major-like and minor-like introns showed an increase in median SI_ES_ relative to their major and minor counterparts, mirroring the differences observed for SI_RET_. Non-canonical introns continued to exhibit high levels of exon skipping, which surpassed the other classes. Despite their low numbers, hybrid introns had statistically higher levels of exon skipping than minor introns. Moreover, the median level of exon skipping for minor introns was statistically lower than that of major introns **(Fig. 4B, Dataset S11 and S12)**. These findings reveal that despite occupying similar spatial locations, rare intron classes exhibit distinct patterns of intron retention and alternative splicing, indicating class-specific splicing regulation beyond 3D positioning.

**Figure 4.**
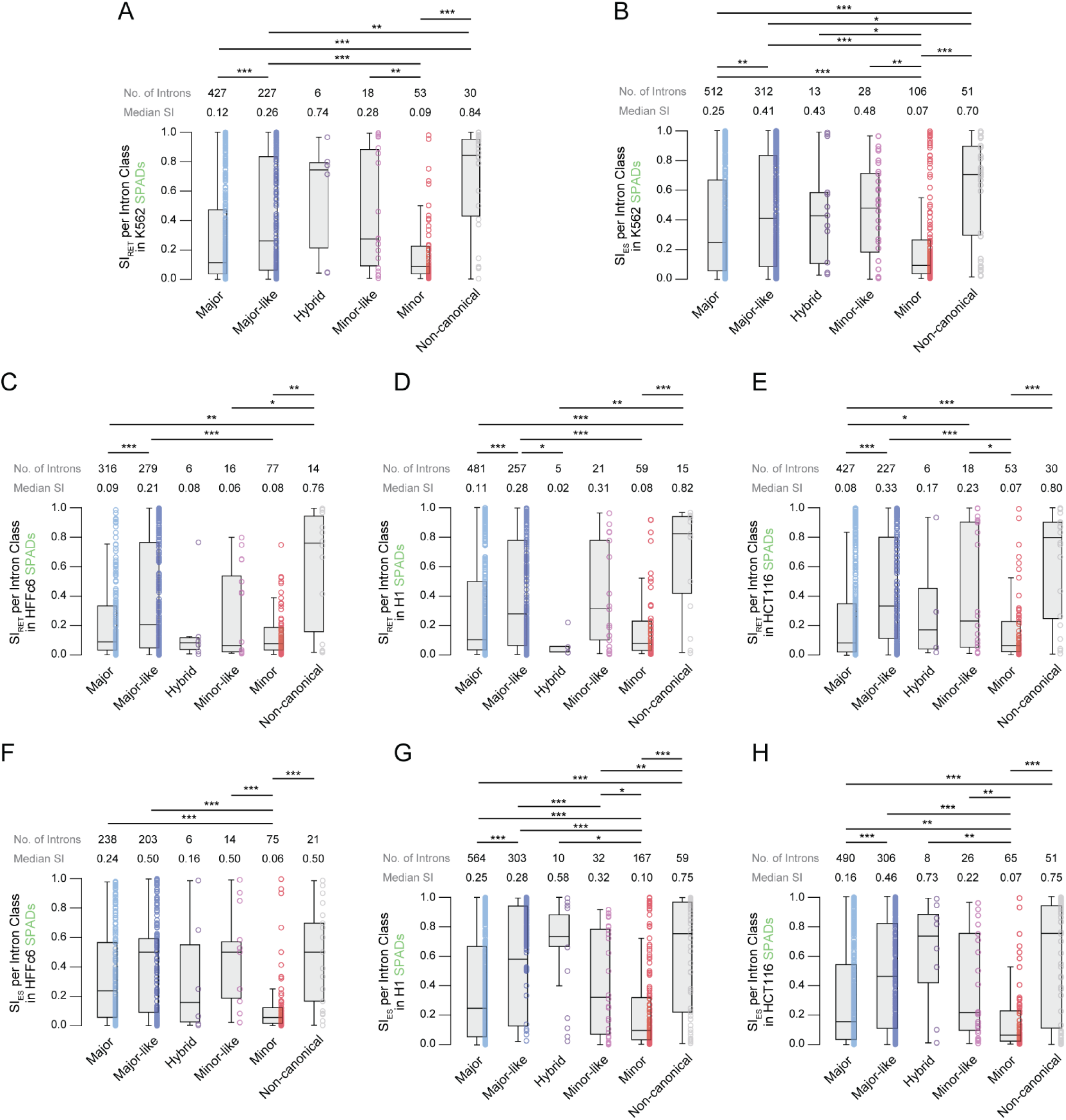
Splicing of rare intron classes in SPADs is conserved across cell lines. A-B. Boxplots depicting the distribution of intron retention (A) and exon skipping (B) in SPADs of K562 cells for each intron class. C-E. Boxplots depicting the distribution of intron retention values for each intron class in SPADs of HFFc6 (C), H1 (D), and HCT116 (E) cells. F-G. Boxplots depicting the distribution of exon skipping values for each intron class in SPADs of HFFc6 (F), H1 (G), and HCT116 (H) cells. \**P*<0.05, \*\**P*<0.01, \*\*\**P*<0.001.

### Cell type-specific splicing of rare intron subclasses in SPADs

Given that K562 cells are derived from chronic myeloid leukemia and MIGs are known to support cancer progression, we next asked whether the observed differences in splicing efficiency were cell type-specific (81). To this end, we analyzed TSA-seq and RNA-seq datasets from three additional cell lines: non-cancerous fibroblasts (HFFc6), embryonic stem cells (H1), and metastatic colorectal cancer cells (HCT116) (3, 79, 80). Data from HFFc6 cells showed increased intron retention of major-like introns compared to both major and minor introns. Notably, we found no significant difference between minor and minor-like intron retention. Non-canonical introns had statistically higher levels of intron retention compared to major, minor-like, and minor introns **(Fig. 4C)**. Like HFFc6 cells, major-like introns exhibited more intron retention in SPADs of H1 cells compared to major, minor, and, surprisingly, hybrid introns **(Fig. 4D)**. Once again, non-canonical introns showed higher levels of intron retention compared to minor, hybrid, and major introns in SPADs **(Fig. 4D)**. The trend of major-like introns exhibiting more intron retention compared to major and minor introns continued in HCT116 cells, except minor-like introns now showed higher levels of intron retention compared to minor introns, like that which was observed in K562 cells **(Fig. 4E)**. Minor-like introns were also retained at higher levels than major introns in SPADs of HCT116 cells **(Fig. 4E)**.

For exon skipping, interrogation of additional cell lines revealed overall conservation of splicing trends observed in K562 cells with some variation. For example, minor introns in SPADs underwent significantly less exon skipping in all three cell lines compared to major introns in SPADs **(Fig. 4F-H)**. While minor-like introns showed higher levels of exon skipping than minor introns in every cell line, major-like introns showed a significant increase in exon skipping compared to major introns in every cell line except HFFc6 **(Fig. 4F-H)**. Non-canonical introns consistently demonstrated elevated levels of exon skipping in every cell line when compared to minor introns but not major introns **(Fig. 4F-H)**. Notably, minor introns in SPADs demonstrated significantly lower levels of exon skipping compared to every other intron class in K562, H1, and HCT116 cells but not HFFc6 cells, where minor introns exhibited significantly less exon skipping compared to every class except hybrid **(Fig. 4B**, **Fig. 4F-H, Dataset S12)**. This finding aligns with the observation that minor introns undergo less exon skipping overall than major introns **(Table S4)** and suggests that their splicing within SPADs is consistently efficient across diverse cell lines.

The dynamic splicing efficiency of rare intron classes within SPADs prompted us to explore how splicing within SPADs varies by intron class across cell lines. For instance, major introns in SPADs of H1 and K562 cells showed elevated intron retention compared to those in SPADs of HCT116 cells **(Fig. 5A)**. This pattern did not hold for minor introns in SPADs, which showed similar splicing across cell lines **(Fig. 5B)**. However, direct comparison is complicated by the fact that SPADs of each cell line consist of a distinct set of introns. Thus, we identified a subset of 173 major introns found in SPADs across all four cell lines and compared their splicing **(Fig. 5C)**. Only one significant comparison was observed, major introns shared across SPADs were more efficiently spliced in HCT116 cells than K562 cells. A similar analysis of minor introns revealed 11 introns in SPADs across all four cell lines with no significant differences in intron retention **(Fig. 5D)**.

**Figure 5.**
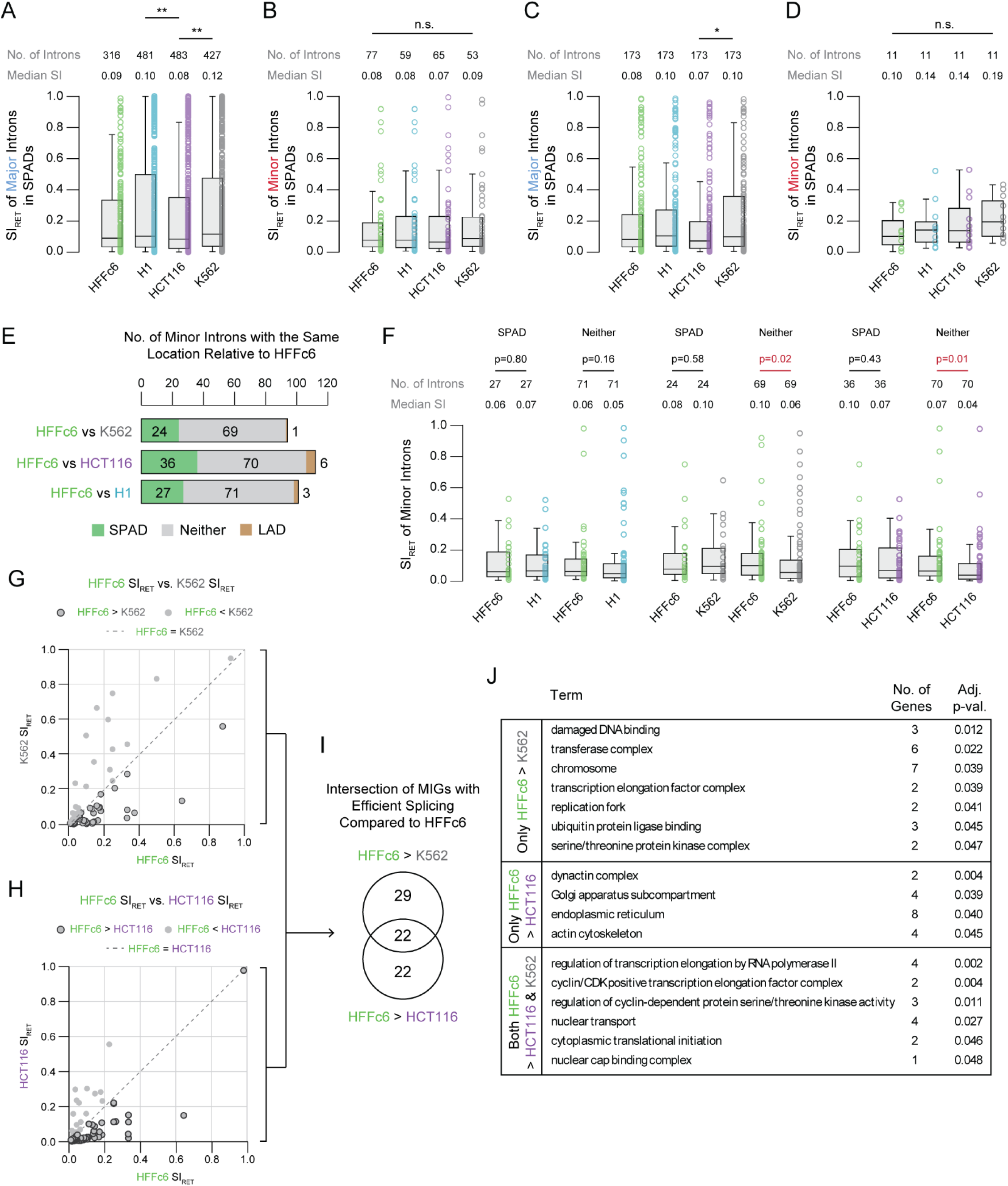
Minor introns outside of SPADs are spliced more efficiently in K562 and HCT116 cancer cells than in HFFc6 fibroblast cells. A-B. Boxplots depicting the distribution of intron retention values for all major (A) and minor (B) introns that pass filtering in each cell line. C-D. Boxplots depicting the distribution of intron retention values for 173 major introns (C) and 11 minor introns (D) with sufficient data across all cell lines. E. The number of introns with the same nuclear body associated location (SPAD, LAD, or “neither”) in HFFc6 and K562, HCT116, or H1 cells in pairwise comparisons. F. Boxplots depicting the distribution of intron retention values for introns with the same location in HFFc6 cells and either H1, K562, or HCT116 cells. G-H. Scatter plots of SI_RET_ values of MIGs in HFFc6 cells and K562 (G) or HCT116 (H) cells. I. Intersection of MIGs with efficient splicing in either K562 and/or HCT116 cells relative to HFFc6 cells. J. GO terms for MIGs with only more efficient splicing in K562 relative to HFFc6, only more efficient splicing in HCT116 relative to HFFc6, or more efficient splicing in both K562 and HCT116 relative to HFFc6. \**P*<0.05, \*\**P*<0.01, \*\*\**P*<0.001.

### Efficient splicing of minor introns in cancer cells is independent of their localization within SPADs

Amongst the four cell lines analyzed, HFFc6 and H1 are both non-cancerous cell lines (3). Given the known role of the minor spliceosome in regulating cell proliferation (37, 68, 81, 93, 94), we chose to use the HFFc6 cell line as a reference to further dissect how spatial context influences minor intron splicing. For each intron found in SPADs of HFFc6, we asked if that same intron was found in SPADs of K562, HCT116, and H1 cells **(Fig. 5E)**. For example, of the 94 minor introns found in SPADs of HFFc6 cells, we found 24 were also in SPADs of K562 cells. Similarly, 36 minor introns were in SPAD of both HFFc6 and HCT116 cells, while 27 minor introns were in SPADs of both HFFc6 and H1 cells. We also tracked overlap between minor introns in LADs and the “neither” category between HFFc6 and the other cell lines. Only one minor intron was found in LADs of both K562 and HFFc6 while 69 minor introns were categorized as “neither” in both HFFc6 and K562 **(Fig. 5E)**. 70 minor introns were in the “neither” category and 6 were in LADs of both HFFc6 and HCT116 cells (**Fig. 5E**). Likewise, 71 were in the “neither” category and 3 were in LADs of both HFFc6 and H1 cells (**Fig. 5E**). We next asked whether minor introns found in the same location across comparisons had similar levels of intron retention. The 27 minor introns in SPADs of both HFFc6 and H1 cells showed no significant differences in splicing, nor did the 71 introns in the “neither” category **(Fig. 5F)**. Similarly, the 24 and 36 minor introns shared in SPADs of HFFc6 to K562 and HCT116, respectively, also exhibited no difference in splicing efficiency. However, minor introns in the “neither” category in both K562 and HCT116 showed lower levels of intron retention compared to the same introns in HFFc6 cells **(Fig. 5F)**. This suggests that minor intron splicing in a non-SPAD context is enhanced in cancer cells relative to non-cancerous fibroblasts.

This 3D context-dependent decrease in intron retention of minor introns found in the “neither” category of K562 and HCT116 cells relative to HFFc6 cells prompted us to identify the introns driving this difference. Therefore, we compared individual SI_RET_ values between HFFc6 and each cancer cell line, identifying minor introns where SI_RET_ was greater in HFFc6 **(Fig. 5G & H)**. We then intersected these intron lists to understand if the same introns were driving the decrease in intron retention in both K562 and HCT116 cells. This revealed 32 minor introns (29 parent genes) with lower intron retention in K562, 33 minor introns (22 parent genes) with lower intron retention in HCT116, and 38 minor introns (22 parent genes) with lower intron retention in both cell lines relative to HFFc6 cells **(Fig. 5I)**. We hypothesized that these differences may reflect cell type-specific and shared biological requirements. Indeed, GO term enrichment analysis of the 29 K562-specific MIGs revealed terms such as damaged DNA binding, chromosome, transcription elongation factor complex, and ubiquitin protein ligase binding **(Fig. 5I and J, Dataset S13)**. The 22 HCT116-specific MIGs were enriched for dynactin complex, endoplasmic reticulum, and actin cytoskeleton. Notably, the 22 MIGs shared between both cancer cell lines were enriched for transcription elongation, cyclin/CDK positive transcription elongation factor complex, and nuclear transport **(Fig. 5I and J)**. These findings suggest that cancer cells enhance minor intron splicing to support gene networks involved in proliferation, stress response, and survival. Importantly, this enhanced splicing is not dependent on proximity to nuclear speckles, suggesting increased expression of minor spliceosome components during oncogenesis overcomes spatial constraints and enables efficient splicing throughout the nucleus.

## Discussion

Proximity of introns to 3D structures such as nuclear speckles, has been shown to improve splicing efficiency and gene expression (3, 4, 32, 61–63). Unlike previous studies that treated all introns as one group, our analysis explores how 3D location informs the splicing of rare intron subclasses (49). While major and minor introns are removed by the major and minor spliceosomes, respectively, the splicing of the remaining intron classes is unclear and likely involves components of one or both spliceosomes (50, 98). We reasoned the 3D distribution of these rare introns may offer insight into their regulation. For example, one could imagine that minor introns located on different chromosomes may cluster in 3D space, creating hubs that facilitate their interaction with lowly abundant minor spliceosome components. Indeed, minor introns are highly enriched in compartment A and SPADs **(Fig. 2B**, **Fig. 3B)**. In contrast, we found that minor-like introns, which are highly similar to minor introns but have acquired features that overlap with major introns such as a -1 G and a weak poly pyrimidine track (49), behave more like major introns in their 3D localization **(Fig. 2B**, **Fig. 3A)**. We have previously proposed that minor-like introns represent a transitory node of flux from minor to major introns and that they might be using an intermediate mechanism of splicing (49). Thus, while their sequence features suggest reliance on the minor spliceosome for removal, the spatial distribution of minor-like introns suggests they do not occupy the same strategic positions as minor introns. Furthermore, hybrid and non-canonical introns are underrepresented in transcriptionally active regions, likely because their sequence features, which lack similarity to major and/or minor introns, restrict their association with 3D regulatory features.

With respect to 3D location and splicing, it was surprising that efficient splicing observed for major and minor introns in SPADs did not extend to major-like or minor-like introns **(Fig. 4)**. This finding suggests that sequence deviation in major-like and minor-like introns is sufficient to decouple splicing efficiency from 3D positioning, specifically in SPADs **(Fig. 4)**. The same phenomenon for non-canonical introns likely reflects their lack of similarity to either major or minor introns, suggesting that they require a complex splicing mechanism. We noted that efficient splicing of rare introns in SPADs is inversely related to alternative splicing (exon-skipping), which is in keeping with previous models linking splicing efficiency with alternative splicing outcomes (68, 95, 99, 100). However, minor introns deviated from this trend, maintaining consistently efficient splicing across diverse cellular and 3D contexts, underscoring their unique regulatory properties.

Minor introns consistently exhibited efficiently splicing within SPADs across cell lines **(Fig. 4**, **Fig. 5D)**. This finding agrees with the crucial role of MIGs in cell division, DNA repair, snRNA biogenesis, and transcription (37, 68). However, when HFFc6, a normal fibroblast cell line, was used as a reference, we found that minor introns outside SPADs were more efficiently spliced in cancer cells **(Fig. 5F)**. This finding aligns with the known role of MIGs in executing oncogenic programs and our previous report linking efficient minor intron splicing to metastatic progression (81, 101, 102). In light of our recent finding that minor spliceosome components are upregulated in cancer cells, these results suggest that enhanced minor intron splicing in cancer cells is driven by elevated minor spliceosome activity rather than spatial positioning. In all, this study provides the first integrative view of 3D genome organization and intron identity, revealing that although rare intron classes occupy distinct spatial locations, splicing efficiency is primarily determined by intron class. These findings establish a framework for future studies on class-specific splicing regulation and its potential roles in development and disease.

## Materials and Methods

### Intron classification

Classification of introns along a spectrum was leveraged from our previously published approach (49), which used position weight matrices (PWMs) to bin introns based on splicing motifs. Coordinates for human introns (hg38, Ensembl v99) from each class were obtained from the prior dataset and used throughout this study. Basic analysis of intron distribution across the linear genome including visualization, quantification of inter-intron distance, and calculation of intron density are described in detail in the supplementary methods.

### Hi-C and other datasets related to 3D genome organization

K562 Hi-C data was compiled from the 4D Nucleome (4DN) Consortium for boundaries and compartments (4DNFI4EFYN3Q and 4DNFIWUAO2QI) **(Dataset S2)** (2). Raw sequencing reads for K562 Hi-C were obtained from GSE63525, processed according to the established 4DN Hi-C Processing Pipeline (https://data.4dnucleome.org/resources/data-analysis/hi_c-processing-pipeline), and used to annotate for TADs and loops as previously described **(Dataset S3-S4)** (103). Annotations for Hi-C subcompartments from K562 were obtained from Xiong and Ma. 2019 (8) through the associated GitHub repository (https://github.com/ma-compbio/SNIPER). SPAD annotations were obtained from 4DNFI625PP2A, 4DNFI6FTPH5V, 4DNFIVZSO9RI, and 4DNFIBY8G6RZ; while LAD coordinates were obtained from 4DNFIP6N54B3, 4DNFIUIDLJJI, 4DNFIV776O7C, and 4DNFICCV71TZ (5). Further details regarding Hi-C and other 3D genome datasets can be found in the supplementary methods.

### Intersection of 3D genomic features with intron coordinates

pybedtools wrapped bedtools intersect was leveraged to assess overlaps between introns and each feature of 3D genome organization (104–106). At least 50% of the intron was required to overlap the 3D genomic feature to be included using the -f option in bedtools intersect set to 0.5. Intersects were then converted into true or false IDs for the presence of each intron in each 3D genomic feature **(Dataset S5 and S6)**. The data was processed to calculate the proportion of introns from each class present in 3D features at various scales, including: compartments, subcompartments, nuclear bodies, and 3D genomic features. Statistical significance was determined with Fisher’s exact test corrected for multiple comparisons with the Bonferroni correction. Details regarding intersection of 3D genomic features with intron coordinates and control bootstrapping and pollution analyses can be found in the supplementary methods.

### RNA sequencing datasets and analysis

RNA sequencing datasets from four human cell lines were received from NCBI Gene Expression Omnibus (GEO) under the following accession numbers per cell line: K562 (GSM958729), H1 (GSM958733), HFFc6 (GSM2687371, GSM2687379), and HCT116 (GSM958749) (79, 80).

A full list of RNA-seq replicates used in this study with SRR numbers can be found in Dataset S8. Fastq files for paired-end read data from H1, HFFc6, K562, and HCT116 cell lines were retrieved using fastq-dump (sratoolkit v2.11.0) with the --gzip –split-files followed by SRR numbers corresponding to the SRA data listed in Dataset S8. Fastq file quality was assessed using fastqc (v0.11.7) and the resulting html files were compiled and visualized using MultiQC (v1.9) (107). Paired-end reads were mapped to hg38 Ensembl v99 using the splice-aware aligner, Hisat2 (v2.2.1) (108). Gene expression was determined by IsoEM2 (v2.0.0) (109), and a gene was considered expressed if the average transcripts per million values (TPM) of its replicates was greater than or equal to one **(Dataset S9)**. This data was then leveraged to filter introns in 3D genomic features into expressed and not expressed categories. Statistical significance was determined with Fisher’s exact test corrected for multiple comparisons with Bonferroni correction.

### Analysis of intron retention

Intron retention analysis was performed using a modified version of our previously published pipeline, which we have employed extensively in prior studies (49, 68, 81, 92, 93, 95). We calculated a splicing index (SI) of intron retention for each intron using BEDTools (v2.29.0) (106). Only introns found in genes expressed at a minimum of 1 TPM, averaged across replicates, that also had at least one supporting read were considered for analysis **(Dataset S9, Fig. S14)**. Introns were then binned into three categories **(Fig. S14 and S15)** and only introns with reads supporting both intron retention and canonical splicing were used for the analyses depicted in Figures 4 and 5. SI values were imported into R (v4.1.0) where subsequent data manipulation and statistical testing was performed using the FSA (v0.9.5) and dunn.test (v1.3.6) packages. Pairwise comparisons were assessed using a Mann-Whitney U test. For multiple group comparisons, we used Kruskal-Wallis test followed by post hoc Dunn’s test with Benjamini-Hochberg correction. Intron retention analysis was performed on 5,000 randomly selected introns for the major and major-like classes. Custom scripts for intron retention analysis are publicly available at https://github.com/amolthof/minor-intron-retention.git

### Alternative splicing analysis

To assess alternative splicing, we again leveraged our previously published bioinformatics pipeline (68). Briefly, uniquely mapped spliced reads spanning each intron of interest were extracted and reads with only one nucleotide mapping to either side of the exon-intron or intron-exon junction were excluded. BEDTools (v2.29.0) was then leveraged to assess exon skipping and cryptic splice site usage (106). A splicing index (SI) value for exon skipping was calculated as described in Fig. S14. Alternative splicing analysis was performed on 5,000 randomly selected introns for the major and major-like classes. Statistical analysis for pairwise and multiple group comparisons was performed using the same approach as described in intron retention analysis.

### Functional enrichment

Gene Ontology (GO) analysis was performed using g:Profiler via its web server interface (https://biit.cs.ut.ee/gprofiler/gost) (110). Multiple correction testing was performed using Benjamini-Hochberg FDR with an adjusted p-value threshold of 0.05. Enriched terms were grouped by biological process, molecular function, and cellular component as described in Dataset S13.

### Code availability

Custom scripts for analysis of intron retention and exon skipping are publicly available at https://github.com/amolthof/minor-intron-retention.git. All other custom scripts used in the current study are publicly available at https://github.com/saren-springer/introns-3Dgenome.

### Data availability

All datasets analyzed in this study are publicly available from NCBI Gene Expression Omnibus, the 4D Nucleome Consortium, and GitHub repository (https://github.com/ma-compbio/SNIPER). Accession numbers for all data files are provided in Datasets S2 and S8.

## Acknowledgements

We thank the Computational Biology Core within the Institute for Systems Genomics at the University of Connecticut for access to the high-performance computing cluster resources. Work in J.E.’s laboratory was supported by the University of Connecticut and an award to J.E. from NIH/NIGMS (R35GM146922). Work in R.N.K.’s laboratory was supported by the University of Connecticut and an award to R.N.K. from Prostate Cancer Foundation (2022 Igor Tulchinksy-Leerom Segal-PCF Challenge Award).

## Author Contributions

Conceptualization – R.N.K. and J.E.; Hi-C methodology – K.F., S.M.R., and J.E.; Intron annotation and RNA-seq methodology – S.M.S., K.N.G., and R.N.K.; investigation – S.M.S., K.F., J.E., and R.N.K.; resources – J.E. and R.N.K.; writing – original draft – S.M.S. and R.N.K.; writing – review and editing – S.M.S., K.F., K.N.G., J.E., and R.N.K.; supervision – J.E. and R.N.K.; funding acquisition – J.E. and R.N.K.

