## Supplemental Information for "Disentangling spatial organization and splicing of rare intron classes in the human genome"

**Contents**

**Supplementary Methods**

Visualizing intron distribution across the linear genome.

Quantification of inter-intron distance.

Calculating intron density across the linear genome.

Hi-C and other datasets related to 3D genome organization.

Intersection of 3D genomic features with intron coordinates.

Bootstrapping and pollution analysis.

Code availability.

**Supplementary Figures**

Figure S1. Characterization of intron abundance per class and chromosome.

Figure S2. Distribution of intron classes along the linear genome based on 250 kb binning.

Figure S3. Statistical analysis of remaining intron classes versus major introns across compartments and subcompartments.

Figure S4. Bootstrapping and permutation analyses to control for class-specific abundance biases when assessing differences in compartment localization.

Figure S5. Bootstrapping and permutation analyses to control for class-specific abundance biases when assessing differences in subcompartment localization.

Figure S6. Statistical analysis of all intron classes versus major introns across 3D genomic features.

Figure S7. Enrichment of genes containing each intron class in the total and core essentialomes.

Figure S8. Compartment distribution of introns in essential and non-essential genes per intron class.

Figure S9. Statistical analysis of remaining intron classes versus major introns in expressed genes across compartments.

Figure S10. Subcompartment enrichment of expressed genes reflects overall compartment biases in the distribution of expressed genes.

Figure S11. Statistical analysis of all intron classes versus major introns in expressed genes across 3D genomic features.

Figure S12. Statistical analysis of remaining intron classes versus major introns in nuclear domains and intersection of introns in compartments and nuclear domains.

Figure S13. Statistical analysis of remaining intron classes versus major introns in expressed genes across nuclear domains.

Figure S14. Bioinformatics pipeline for intron retention and exon skipping analysis.

Figure S15. Quantification of introns in exon skipping and intron retention bins.

**Supplementary Tables**

Table S1. Percentage of the coding genome occupied by major and rare intron classes.

Table S2. Intron abundance per class per chromosome.

Table S3. Results of Fisher exact significance testing (BH adjusted p-values) on proportions of introns per class per 3D genomic feature.

Table S4. Statistical analysis results of Kruskal-Wallis test to assess differences in exon skipping and intron retention between intron classes, regardless of 3D location.

**Supplementary Datasets**

Dataset S1. Intron abundance per 250 kb bin.

Dataset S2. Hi-C and other 3D dataset annotation information per cell line.

Dataset S3. K562 loop anchor annotations.

Dataset S4. K562 TAD annotations.

Dataset S5. Intersects of introns with 3D genomic features in K562, HCT116, HFFc6, and H1 cells requiring 50% intron overlap with each feature.

Dataset S6. Intersects of introns with 3D genomic features in K562 cells requiring 1 bp or 90% intron overlap with each feature.

Dataset S7. List of genes in the total and core essentialomes.

Dataset S8. RNA-seq dataset information per cell line.

Dataset S9. List of expressed genes per cell line.

Dataset S10. Subcompartment expression by intron class.

Dataset S11. Summary of statistical comparison of intron retention across intron classes and cell lines.

Dataset S12. Summary of statistical comparison of exon skipping across intron classes and cell lines.

Dataset S13. GO terms with parent genes of introns with increased splicing efficiency in K562 and HCT116 relative to HFFc6 cells.

**Supplementary References**

**Supplementary Methods**

***Visualizing intron distribution across the linear genome.*** Visualization of intron distribution within the human genome was performed using the chromoMap R package (v0.2.1) (1). Chromosome coordinates were obtained from Ensembl GRCh38 (Ensembl v99). Intron coordinates for each class were previously produced by Olthof-Schwoerer et al. (2). Here, we concatenated annotations from all six classes and generated a 2D genome map using the following parameters: chr.2D.plot = TRUE, n_win.factor = 1, and win.summary.display = FALSE. Intron categories were filtered using the plot_filter argument. Major and major-like introns were excluded from plotting, and the remaining intron classes were displayed using a custom color scheme.

***Quantification of inter-intron distance.*** To calculate the distance between consecutive introns of the same class (i.e. inter-intron distance) BED files for each intron class were parsed using a custom Python script (Python v3.8.1) that leveraged pandas (v1.5.3), glob, and os libraries. Introns were sorted by chromosome and start position prior to calculating the distance between each intron and its nearest neighboring intron of the same class on the same chromosome, excluding overlapping or nested introns. All resulting inter-intron distances were concatenated into a single dataframe and exported for subsequent analysis and plotting.

***Calculating intron density across the linear genome.*** A 250 kb windowing approach was applied to the hg38 reference genome to quantify intron density across the linear genome per intron class. BED-formatted intron coordinates were imported into R (v4.1.0) and parsed for each intron class. Each chromosome was partitioned into 250,000 bp windows and intron count and density (count per 250 kb bin) was calculated. In addition, all possible combinations of intron classes were assessed for co-occurrence within each window. These findings, along with the corresponding gene symbols for each intron per window, were compiled into a genome-wide matrix which can be found in Dataset S1.

***Hi-C and other datasets related to 3D genome organization.*** Raw sequencing reads for K562 Hi-C were obtained from GSE63525, processed according to the established 4DN Hi-C Processing Pipeline (https://data.4dnucleome.org/resources/data-analysis/hi_c-processing-pipeline) with Juicer Tools (v1.22.01) (3), and used to annotate for TADs and loops as previously described, modified using SCALE normalization **(Dataset**

**S3 and S4)** (4, 5). Datasets of boundaries and compartments were obtained from the associated data (4DNFI4EFYN3Q and 4DNFIWUAO2QI) **(Dataset S2)**. The compartment files were split into compartments A and B by assigning positive scores to compartment A and negative scores to compartment B (4DNFIWUAO2QI). Annotations for Hi-C subcompartments from K562 were obtained from Xiong and Ma et al. (6) through the associated GitHub repository (https://github.com/ma-compbio/SNIPER). The genomic coordinates of SPADs from four cell lines (H1, HFFc6, K562, HCT116) were obtained from processing files 4DNFI625PP2A, 4DNFI6FTPH5V, 4DNFIVZSO9RI, and 4DNFIBY8G6RZ as previously described (7). Coordinates of LADs were obtained from 4DNFIP6N54B3, 4DNFIUIDLJJI, 4DNFIV776O7C, and 4DNFICCV71TZ (8). To prevent overcounting any given region of the genome, each dataset was collapsed to merge overlapping intervals as previously described (9). Statistical information about these collapsed final datasets is provided in Dataset S2.

***Intersection of 3D genomic features with intron coordinates.*** To determine the localization of introns from each class in TADs, loop anchors, boundaries, compartments, subcompartments, SPADs, and LADs, overlaps between introns and each feature of 3D genome organization were performed using pybedtools wrapped bedtools intersect (10–12). At least 50% of the intron was required to overlap the 3D genomic feature to be included by using the argument f=0.50. The c=True argument was also used to give the number of times this requirement was satisfied for each intron intersect. The counts were then converted into true or false IDs for the presence of each intron in each inspected 3D genomic feature **(Dataset S5)**. Intersects were also performed with f=0.90, requiring 90% of the intron to be in the 3D genomic feature, and without the f argument, defaulting to 1 bp overlap, to control for requiring differing amounts of the catalytically critical intronic sequence in the overlap **(Dataset S6)**. The data was analyzed to determine the proportion of introns from each class located in features of 3D genome organization at different scales: compartment A, compartment B, or neither; subcompartments A1, A2, B1, B2, and B3; SPAD, LAD, or neither; and TADs, loop anchors, boundaries, combinations of these, or none. Analysis was also conducted to calculate the proportion of introns that overlapped either SPAD or LAD as well as the various compartments or subcompartments. Compartments A and B were examined by summing subcompartments A1 with A2 into “compartment A" and B1 with B2 and B3 into “compartment B”. Statistical significance was determined with Fisher’s exact test corrected for multiple comparisons with the Bonferroni correction.

***Bootstrapping and pollution analysis.*** To assess whether the spatial distribution of rare intron classes within the 3D genome could have arose by chance, we performed a custom bootstrapping and pollution analysis using BEDTools (v2.29.0), BASH scripting, and R scripting (12). For each rare class, we matched the total number of introns in that class (i.e. 850 for minor introns) by randomly sampling an equal number of major introns 1,000 times to generate bootstrapped datasets. To test how the identity of rare introns influences compartmental distribution, we performed a parallel pollution analysis. For each intron class, 1,000 datasets were generated, consisting of between 10% and 100% rare introns (in increments of 10%), with the remainder of introns consisting of major introns. For example, when assessing minor introns (n = 850), 1,000 pollution datasets were generated for each percentage of pollution analysis using a mixture of minor and major introns (e.g. 85 minor + 765 major for 10%), with all introns randomly shuffled. The BED files produced from this analysis were then intersected with a master table of intron annotations in the 3D genome using bedtools intersect and downstream analysis was performed in R. For each bootstrapping and pollution dataset, we calculated the proportion of introns per class that fell into compartments, subcompartments, and nuclear domains. The resulting distribution of proportions was visualized using ggplot2 (13), and Pearson correlation coefficients were calculated for the relationship between rare intron content (0-100%) and the proportion of introns localized to each group (e.g. A, B, or "neither"). Correlation testing was performed independently for each 3D feature, and p-values were corrected for multiple comparisons using Benjamini-Hochberg correction.

***Code availability.*** All custom scripts and code used in the current study are publicly available at our GitHub repository: https://github.com/saren-springer/introns-3Dgenome. This includes code for generating chromoMaps, calculating inter-intron distance, intron density calculations, as well as scripts for bootstrapping and pollution analyses.

### Supplementary Figures

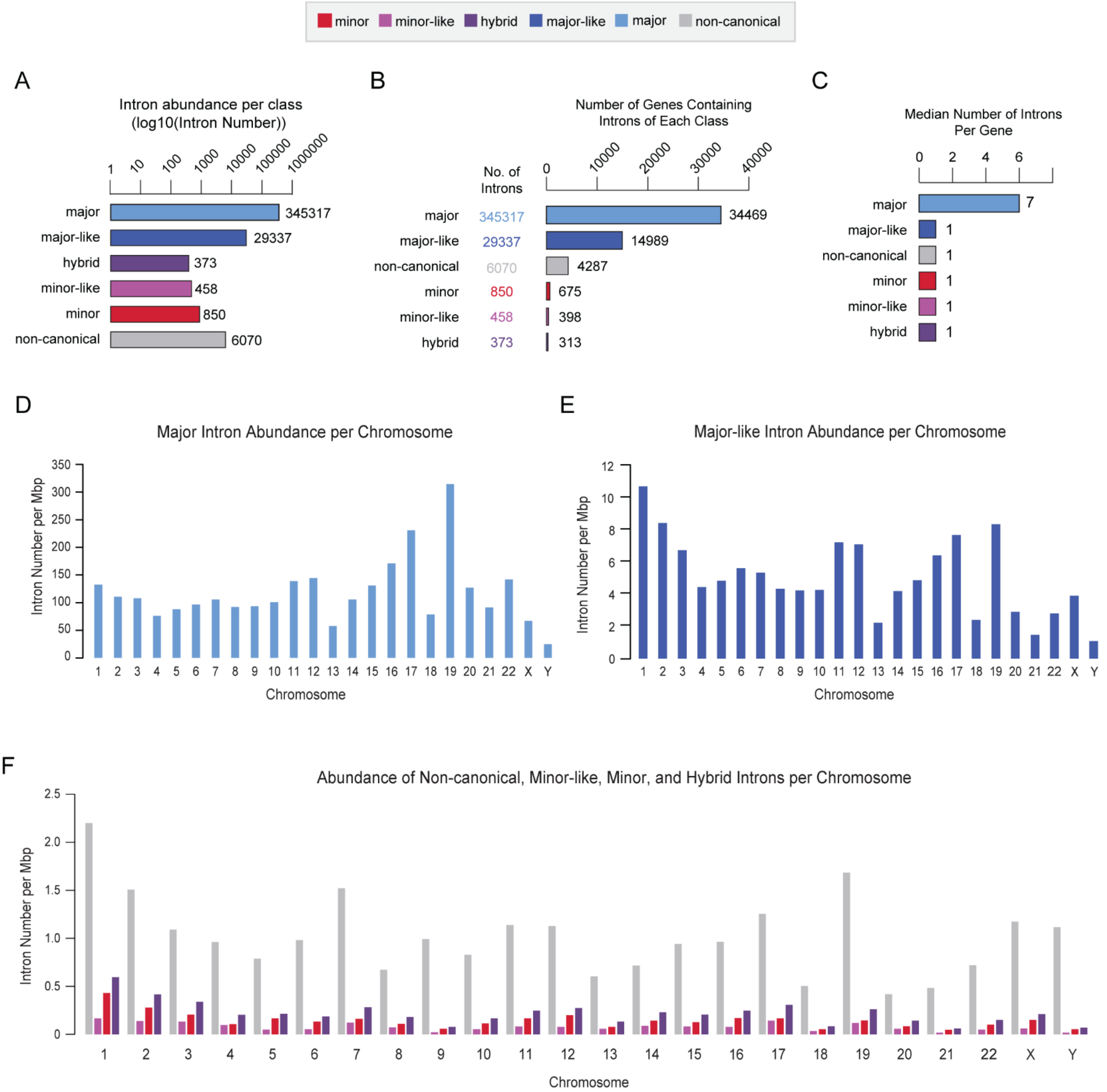


**Figure S1. Characterization of intron abundance per class and chromosome.** A. Bar graph of intron abundance per class presented as log10 transformed values. Actual values are listed to the right of each bar. B. Bar plot of the number of genes that contain introns of each class. C. Bar plot of the median number of introns per gene for each intron class. D-F. Bar graphs of major (D), major-like (E), or non-canonical, minor-like, minor, and hybrid (F) intron abundance per chromosome. Per chromosome count values can be found in Table S2.

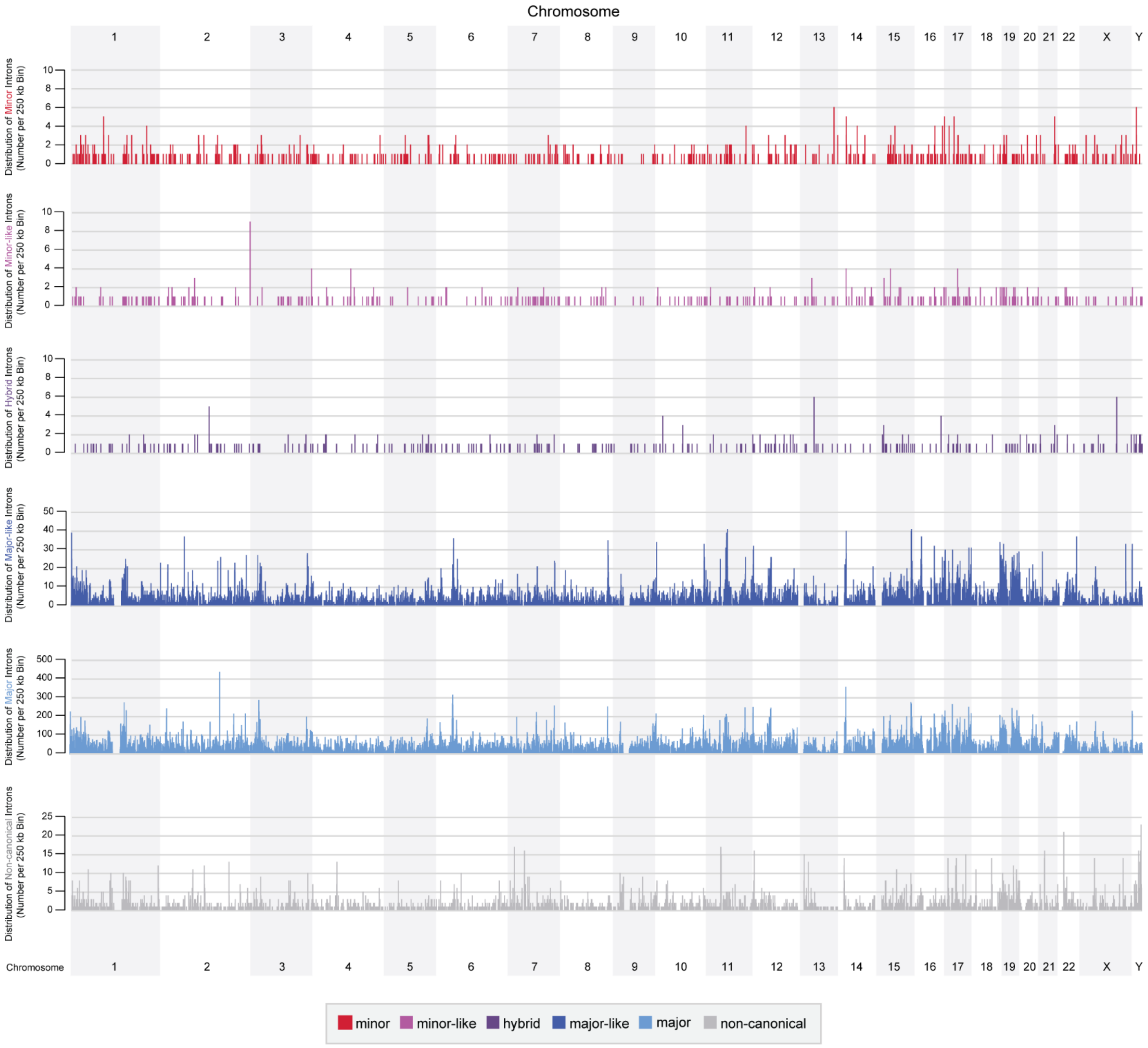


**Figure S2. Distribution of intron classes along the linear genome based on 250 kb binning.** Density plots per intron class. Grey boxes highlight alternating chromosomes.

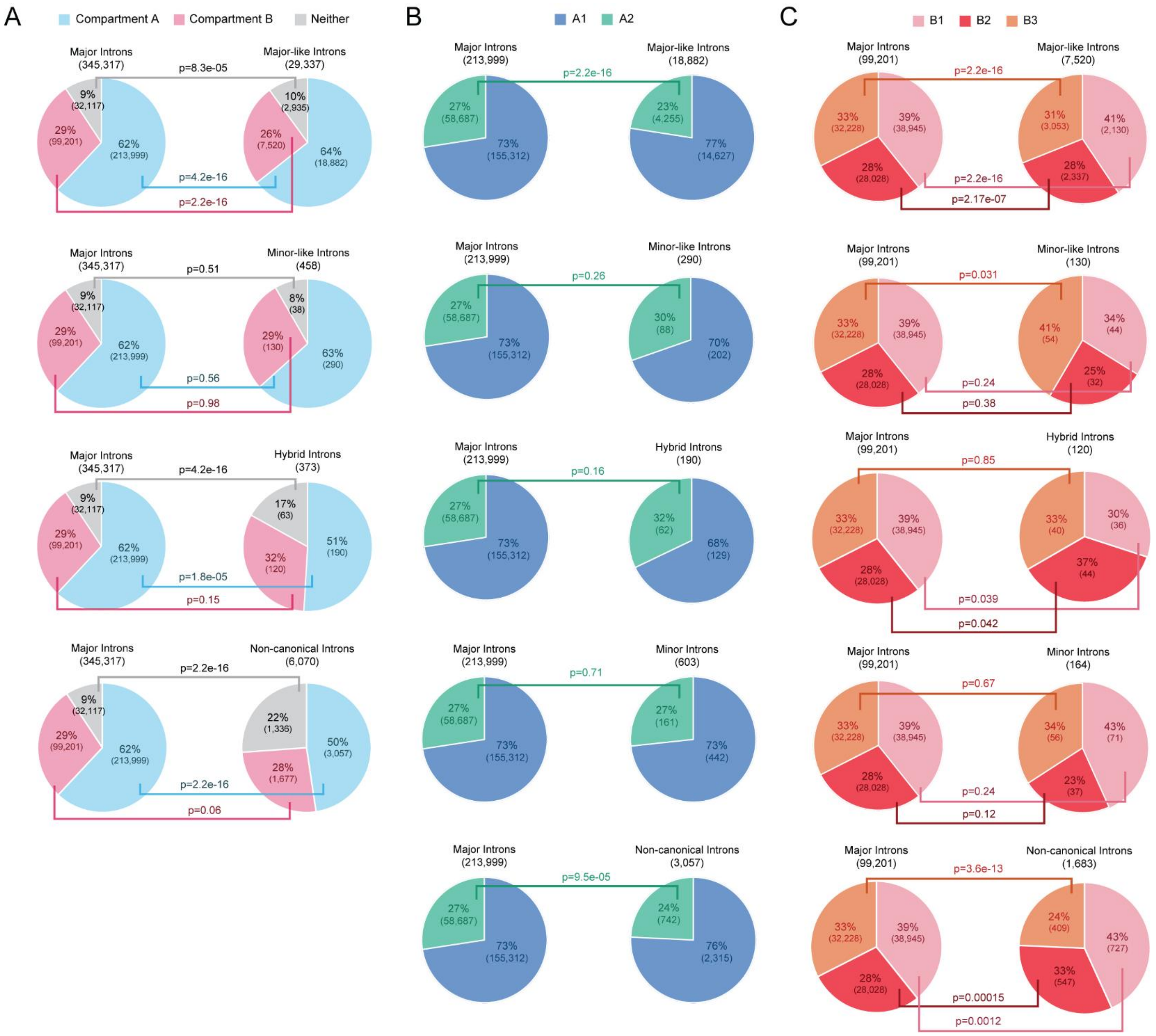


**Figure S3. Statistical analysis of remaining intron classes versus major introns across compartments and subcompartments.** A. Fisher's exact test comparing the compartment distribution of major versus major-like, minor-like, hybrid, and non-canonical introns. B. Fisher's exact test comparing the A1 and A2 subcompartment distribution of major versus all other intron classes. C. Fisher's exact test comparing the B1, B2, and B3 subcompartment distribution of major versus major-like, minor-like, minor, hybrid, and non-canonical introns.

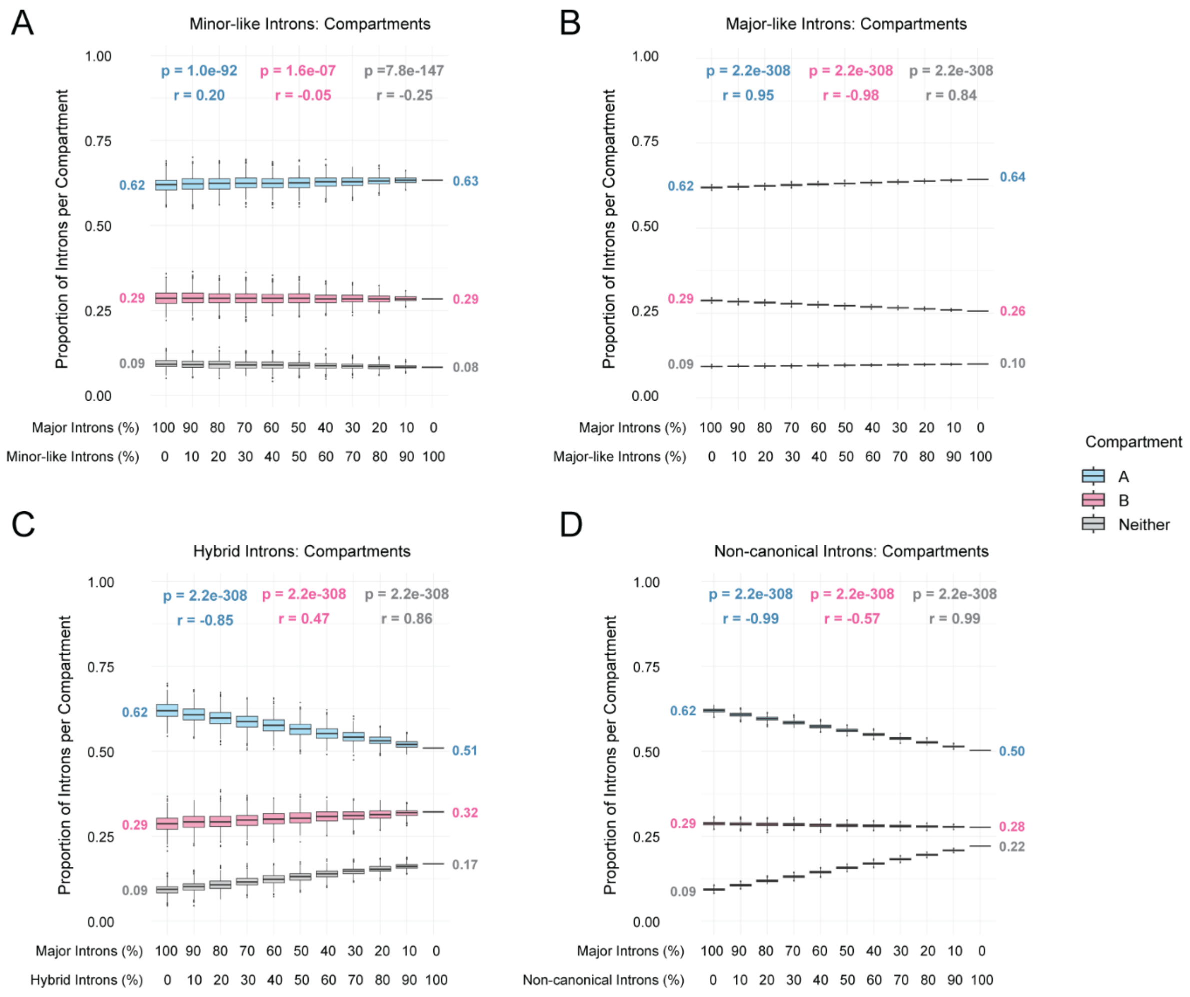


**Figure S4. Bootstrapping and permutation analyses to control for class-specific abundance biases when assessing differences in compartment localization.** A-D. Proportion of introns per compartment across 1000 bootstrapped intron sets that vary the proportion of major introns versus the intron class of interest. Comparisons are shown for minor-like (A), major-like (B), hybrid (C), and non-canonical (D) introns. The first boxplot of each graph represents 1000 permutations of randomly selected major introns, number matched to the total number of introns in the class of interest. The proportion of major introns then progressively decreases across permutations, while the number of introns from the class of interest increases, until the final boxplot, which reflects the observed localization of all introns from that class. Median values for the first and last boxplots are listed on the left and right sides of each graph, respectively. p-values and Pearson’s correlation coefficients (r) are shown at the top of each graph and indicate the significance and strength of the correlation across bootstrapping trials, respectively.

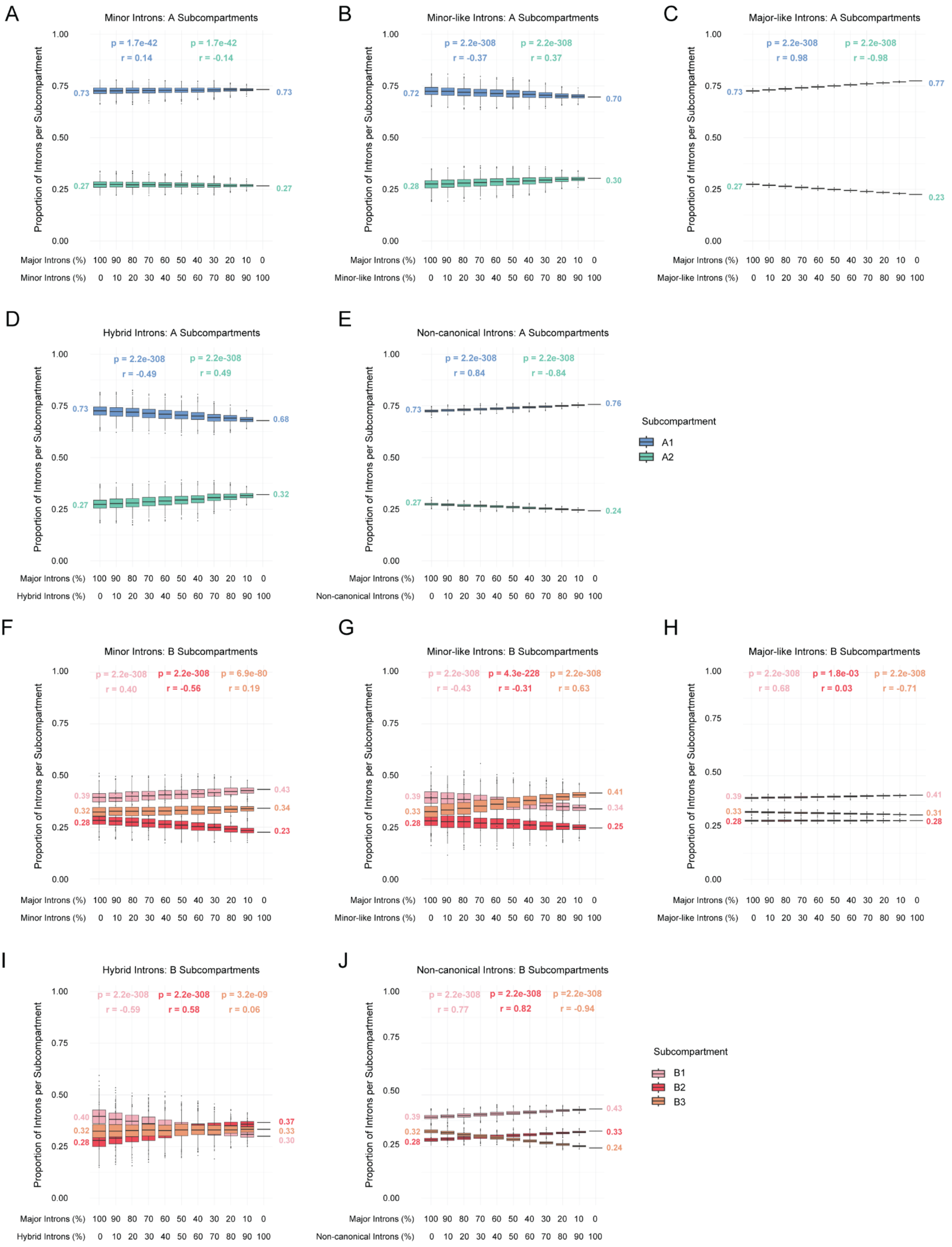
A
Minor Introns: A Subcompartments
p = 1.7e-42
r = 0.14
p = 1.7e-42
r = -0.14
0.73
0.73
0.27
0.27
B
Minor-like Introns: A Subcompartments
p = 2.2e-308
r = -0.37
p = 2.2e-308
r = 0.37
0.72
0.70
0.28
0.30
C
Major-like Introns: A Subcompartments
p = 2.2e-308
r = 0.98
p = 2.2e-308
r = -0.98
0.73
0.77
0.27
0.23
D
Hybrid Introns: A Subcompartments
p = 2.2e-308
r = -0.49
p = 2.2e-308
r = 0.49
0.73
0.68
0.27
0.32
E
Non-canonical Introns: A Subcompartments
p = 2.2e-308
r = 0.84
p = 2.2e-308
r = -0.84
0.73
0.76
0.27
0.24
Subcompartment
A1
A2
F
Minor Introns: B Subcompartments
p = 2.2e-308
r = 0.40
p = 2.2e-308
r = -0.56
p = 6.9e-80
r = 0.19
0.39
0.32
0.28
0.43
0.34
0.23
G
Minor-like Introns: B Subcompartments
p = 2.2e-308
r = -0.43
p = 4.3e-228
r = -0.31
p = 2.2e-308
r = 0.63
0.39
0.33
0.28
0.41
0.34
0.25
H
Major-like Introns: B Subcompartments
p = 2.2e-308
r = 0.68
p = 1.8e-03
r = 0.03
p = 2.2e-308
r = -0.71
0.39
0.33
0.28
0.41
0.31
0.28
I
Hybrid Introns: B Subcompartments
p = 2.2e-308
r = -0.59
p = 2.2e-308
r = 0.58
p = 3.2e-09
r = 0.06
0.40
0.32
0.28
0.37
0.33
0.30
J
Non-canonical Introns: B Subcompartments
p = 2.2e-308
r = 0.77
p = 2.2e-308
r = 0.82
p =2.2e-308
r = -0.94
0.39
0.32
0.28
0.43
0.33
0.24
Subcompartment
B1
B2
B3
Proportion of Introns per Subcompartment
1.00
0.75
0.50
0.25
0.00
Major Introns (%)
100 90 80 70 60 50 40 30 20 10 0
Minor Introns (%)
Minor-like Introns (%)
Major-like Introns (%)
Hybrid Introns (%)
Non-canonical Introns (%)
0 10 20 30 40 50 60 70 80 90 100

**Figure S5. Bootstrapping and permutation analyses to control for class-specific abundance biases when assessing differences in subcompartment localization.** A-E. Proportion of introns per A1 and A2 subcompartments across 1000 bootstrapped intron sets that vary the proportion of major introns versus the intron class of interest. Comparisons are shown for minor (A), minor-like (B), major-like (C), hybrid (D), and non-canonical (E) introns. F-J. Proportion of introns per B1, B2, and B3 subcompartments across 1000 bootstrapped intron sets that vary the proportion of major introns versus the intron class of interest. Comparisons are shown for minor (F), minor-like (G), major-like (H), hybrid (I), and non-canonical (J) introns. Median values for the first and last boxplots are listed on the left and right sides of each graph, respectively. p-values and Pearson's correlation coefficients (r) are shown at the top of each graph and indicate the significance and strength of the correlation across bootstrapping trials, respectively.

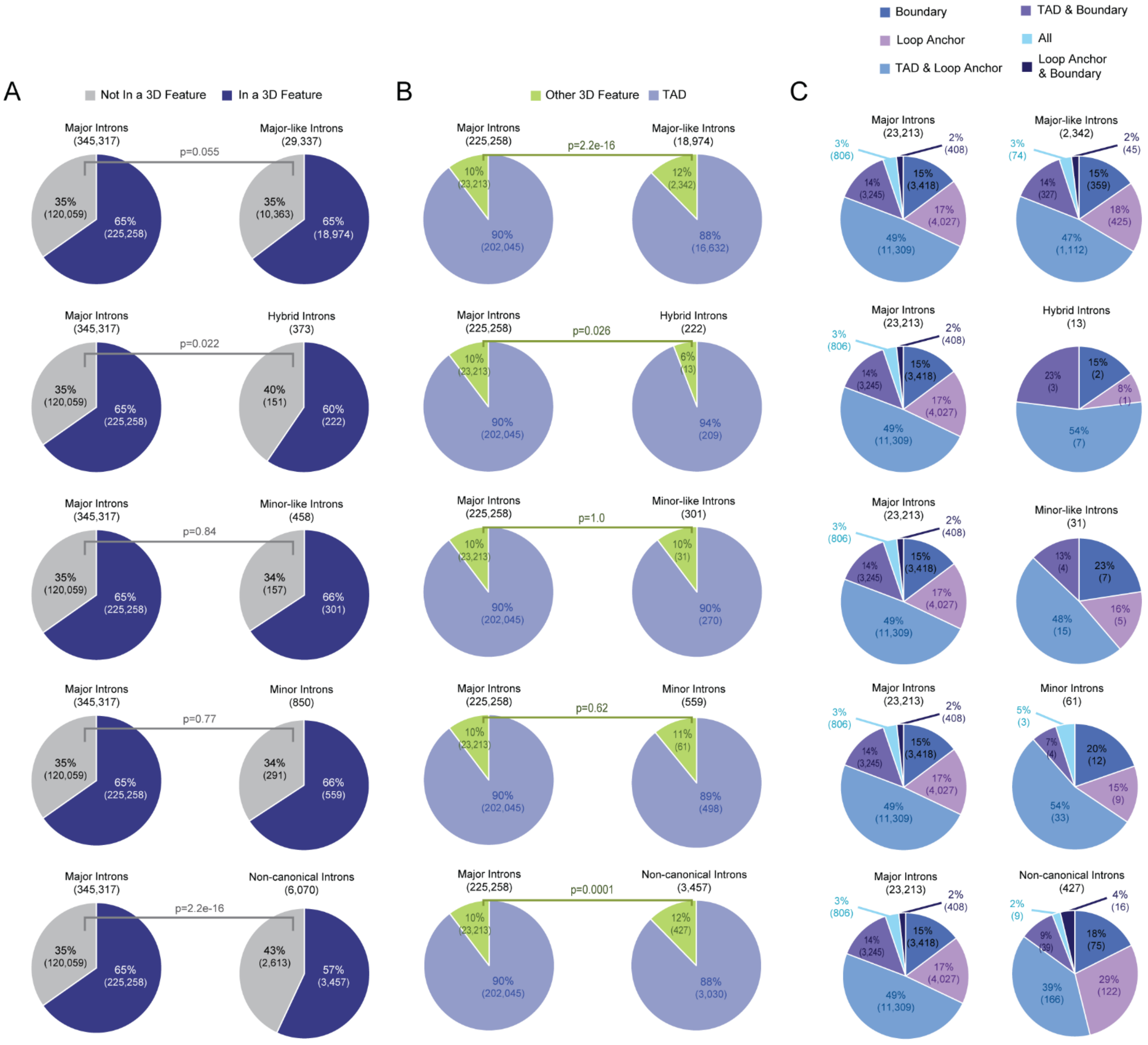


**Figure S6. Statistical analysis of all intron classes versus major introns across 3D genomic features.** A. Fisher's exact test comparing the number of major introns versus all other intron classes that overlap with any 3D genomic feature. B. Fisher's exact test comparing the number of major introns versus all other intron classes that overlap with TADs or another 3D genomic feature. C. Comparison of the number of major introns versus all other intron classes that overlap with any combination of 3D features. Fisher's exact results for panel C can be found in Table S3.

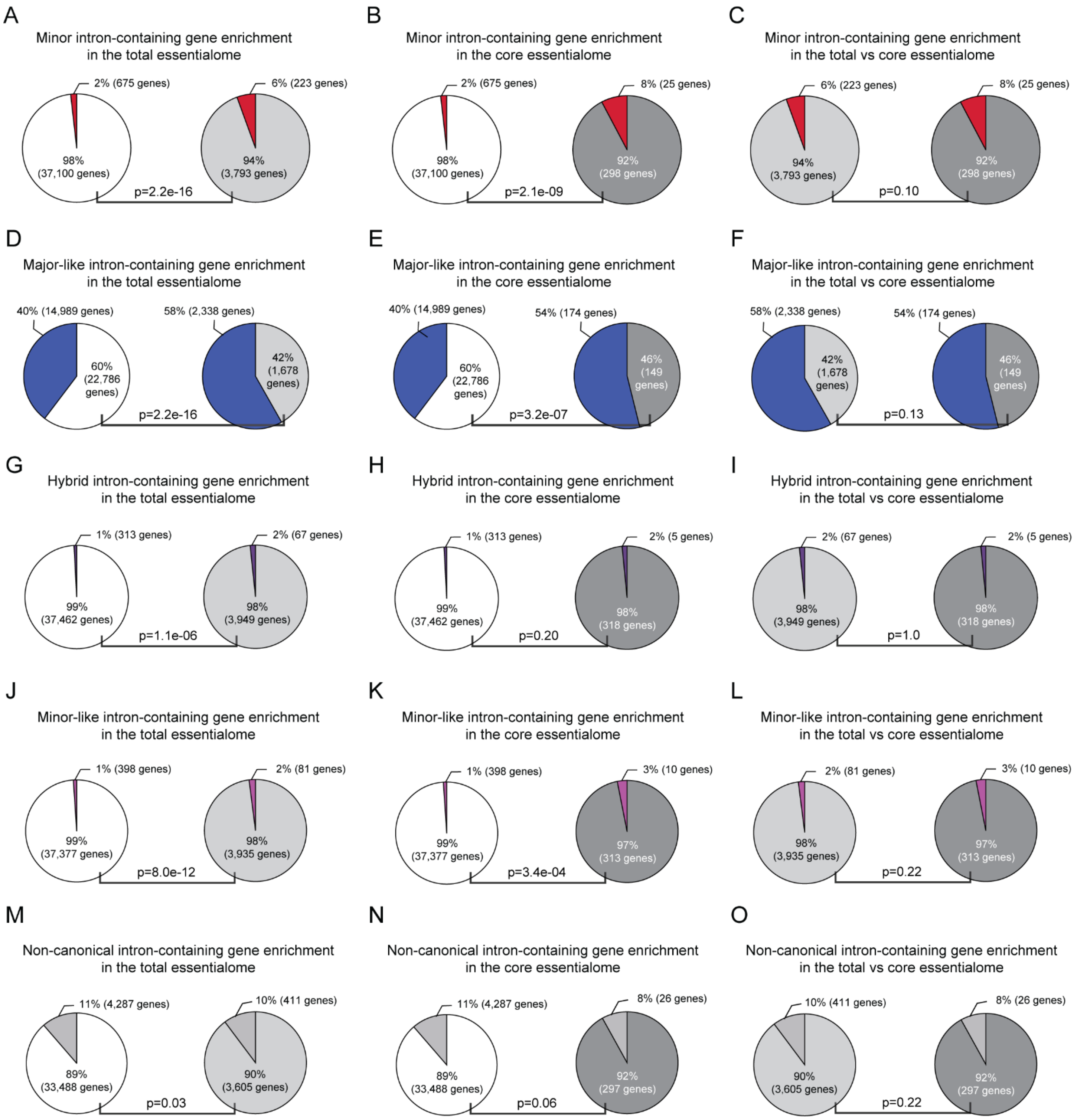


**Figure S7. Enrichment of genes containing each intron class in the total and core essentialomes.** Fisher's exact tests were performed to determine whether genes containing a given intron class were enriched in the total or core essentialome compared to their overall genomic abundance. In each pair of pie charts in panels A, B, D, E, G, H, J, K, M, and N, the left chart is enrichment of all genes containing the intron class of interest relative to all interrogated genes while the right chart is enrichment of essential genes containing the intron class of interest relative to all essential genes. This analysis was performed for genes containing minor (A-C), major-like (D-F), hybrid (G-I), minor-like (J-L), and non-canonical (M-O) introns. Panels C, F, I, L, and O compare enrichment between the total and core essentialome per intron class.

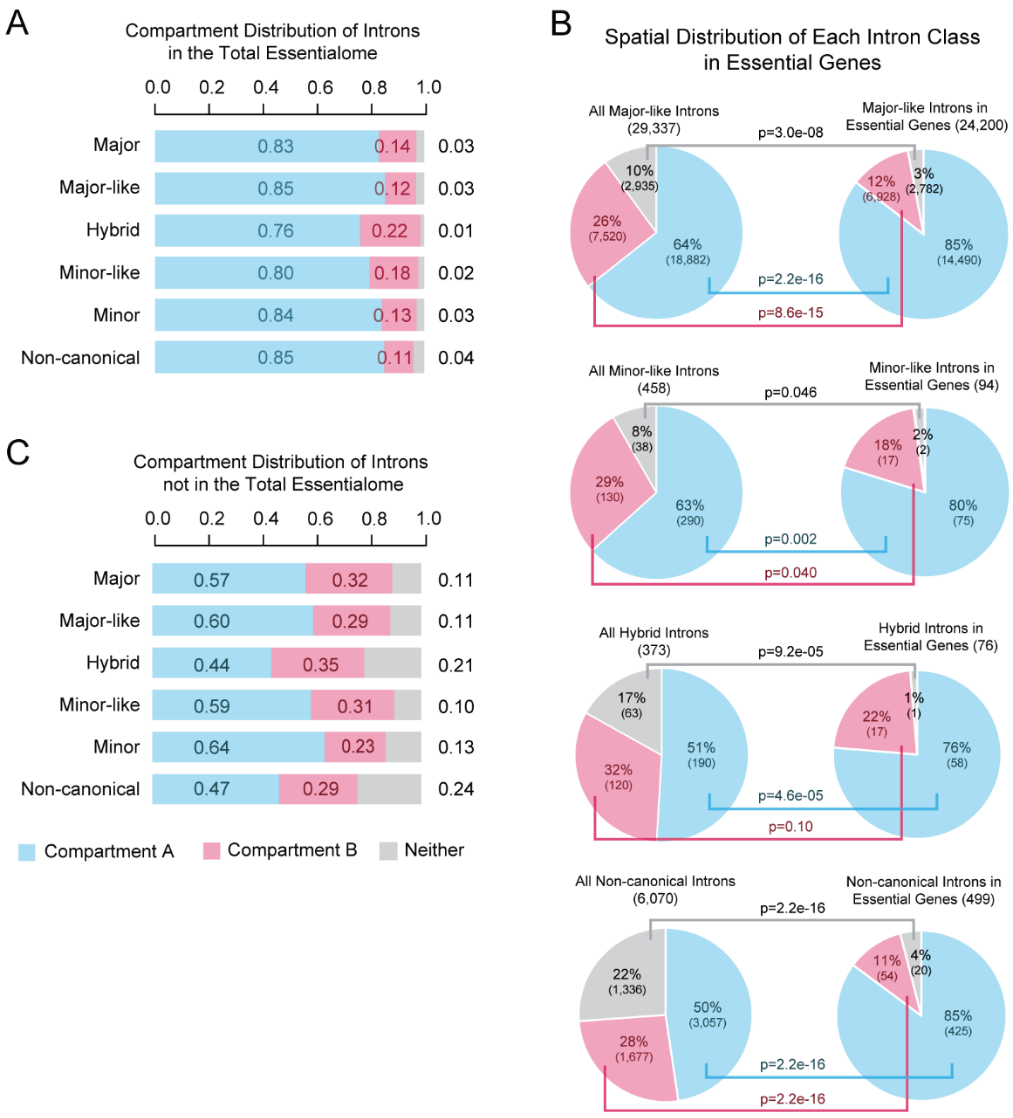


**Figure S8. Compartment distribution of introns in essential and non-essential genes per intron class.** A. Proportional distribution of essential genes containing each intron class across compartments A, B, and neither. B. Fisher's exact testing comparing the spatial distribution of each intron class in essential genes to the overall distribution of all introns in that class. C. Proportional distribution of non-essential genes containing each intron class across compartments A, B, and neither.

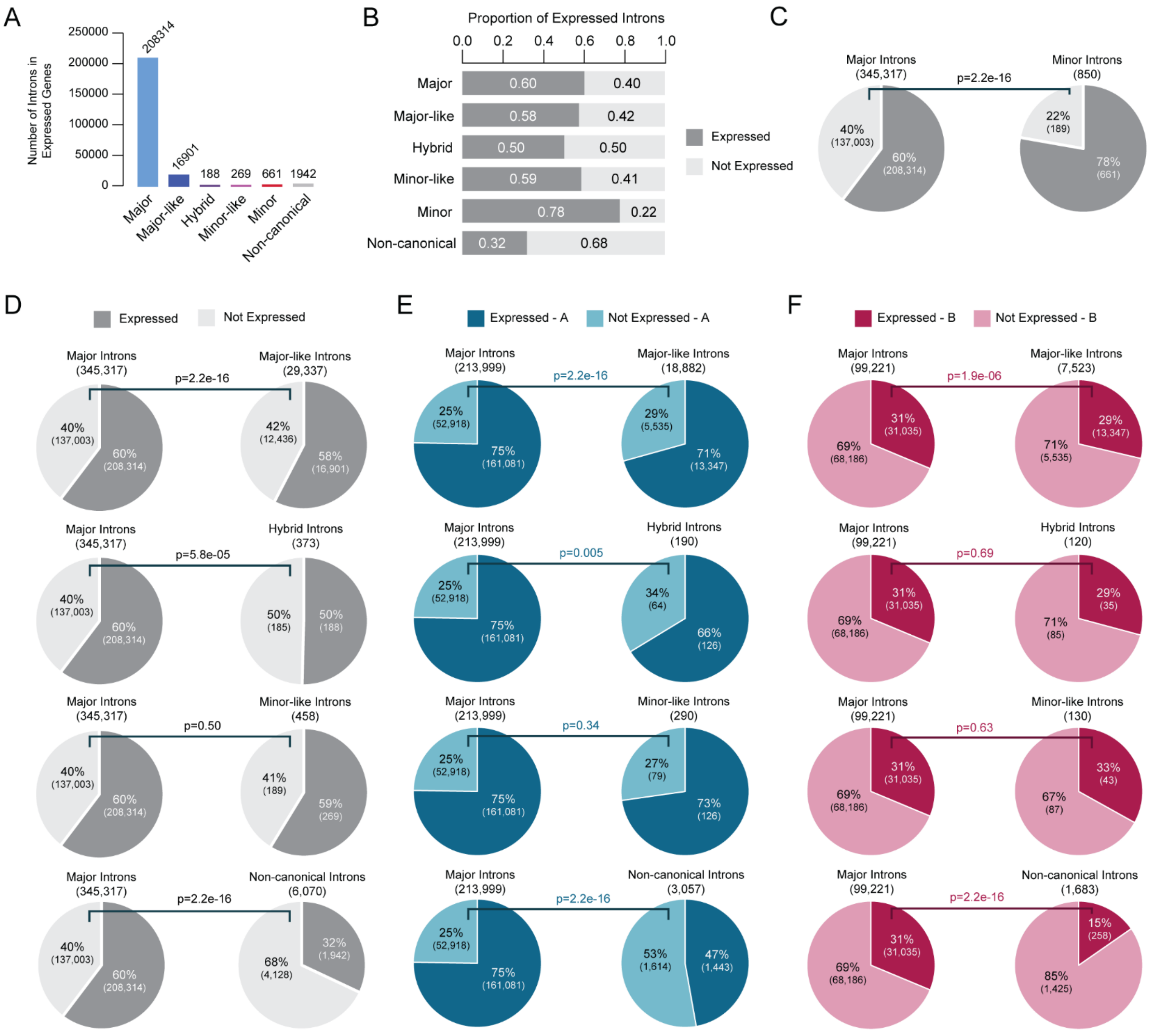


**Figure S9. Statistical analysis of remaining intron classes versus major introns in expressed genes across compartments.** A. Bar plot of the number of introns per class in expressed genes. B. Proportion of introns per class in expressed genes. C. Fisher's exact test comparing the distribution of major versus minor introns in expressed genes. D. Fisher's exact test comparing the distribution of major versus minor-like, major-like, hybrid, and non-canonical introns in expressed genes. E. Fisher's exact test comparing the distribution of major versus all other intron classes in compartment A and in expressed genes. F. Fisher's exact test comparing the distribution of major versus all other intron classes in compartment B and in expressed genes.

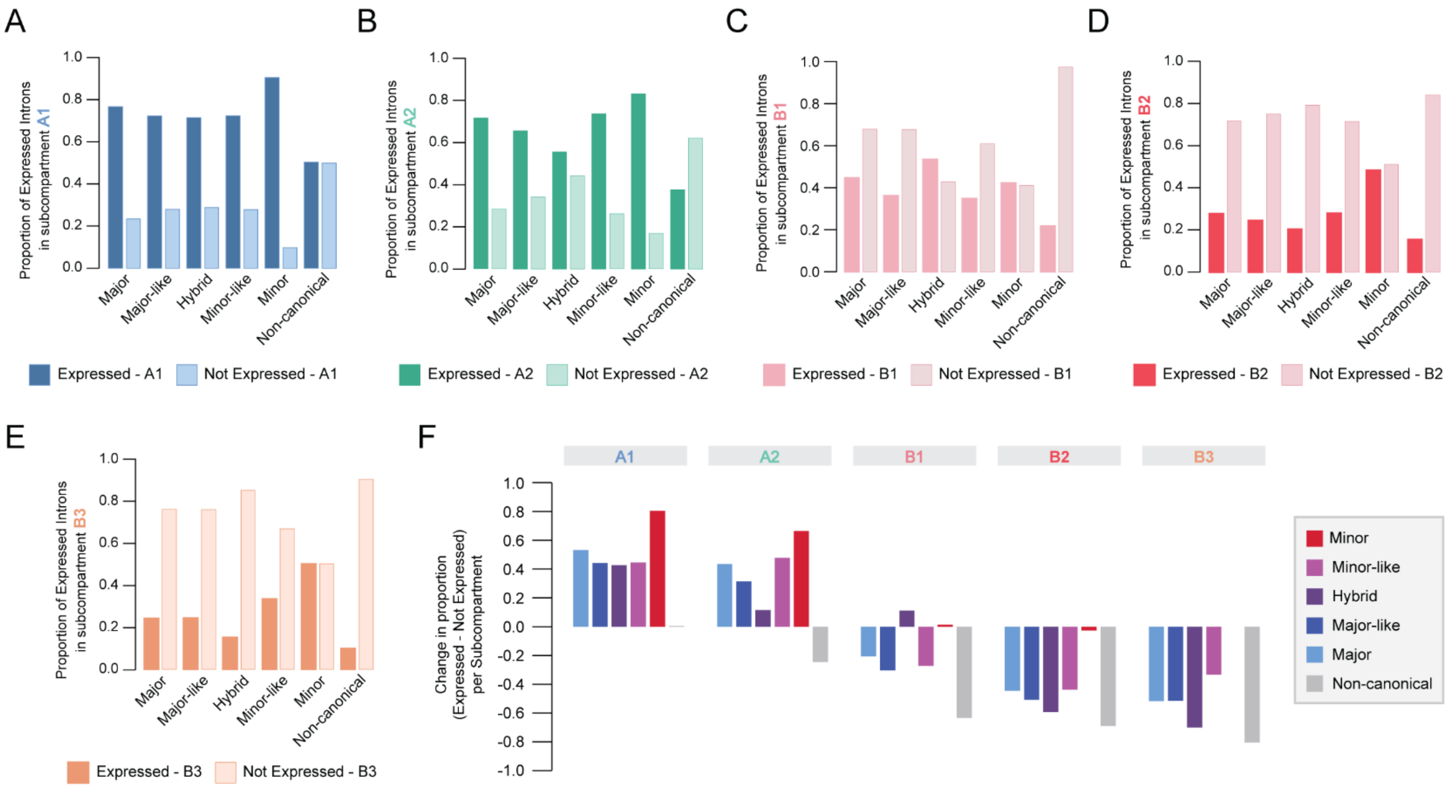


**Figure S10. Subcompartment enrichment of expressed genes reflects overall compartment biases in the distribution of expressed genes.** A-E. Bar charts showing the proportion of introns per class within expressed genes, subset by subcompartments A1, A2, B1, B2, and B3. F. Bar plot of the change in proportion of expressed and not expressed introns per subcompartment. Values for proportions and change in proportions per intron class can be found in Dataset S10.

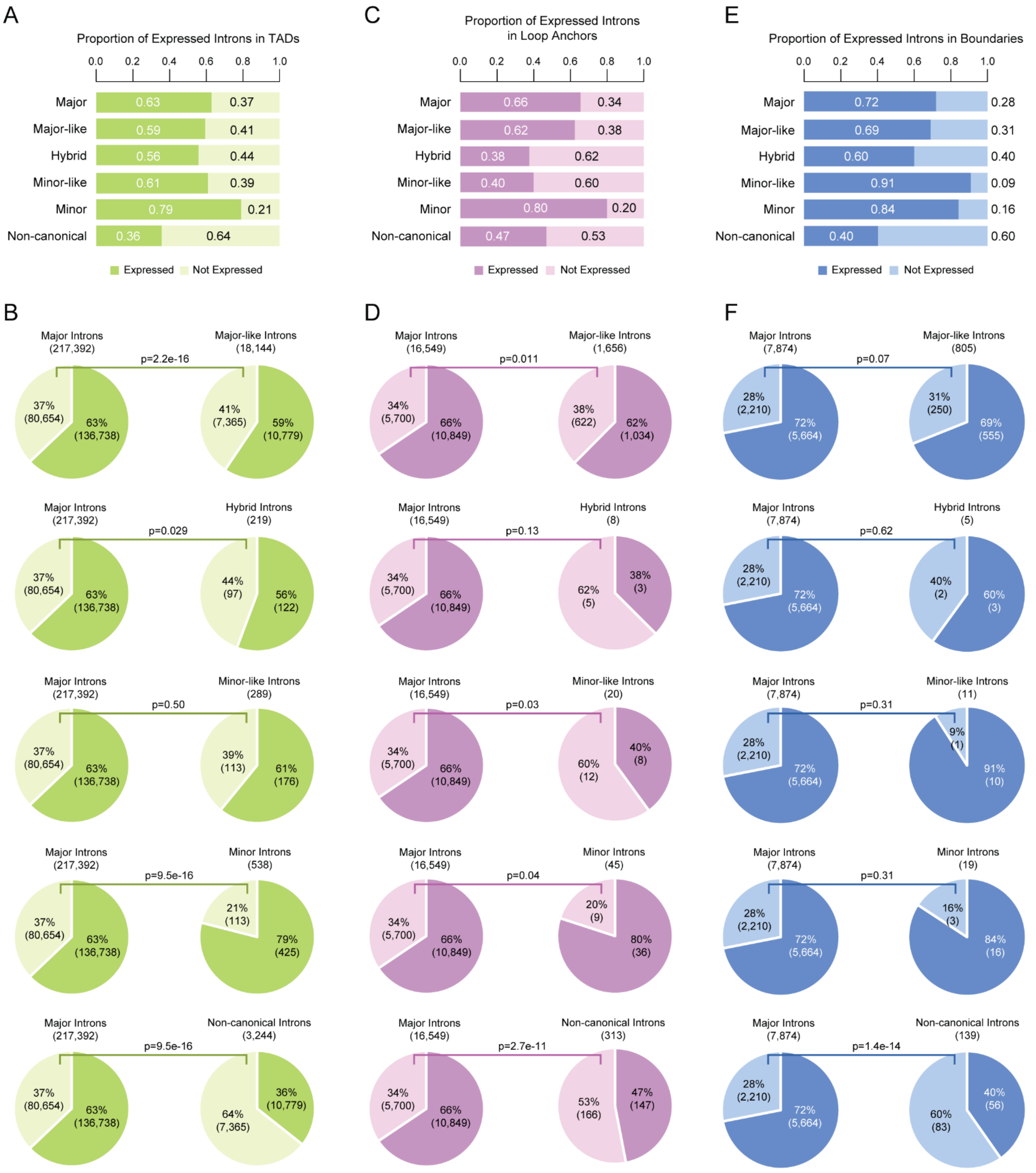


**Figure S11. Statistical analysis of all intron classes versus major introns in expressed genes across 3D genomic features.** Proportion of introns per class that are in TADs (A), Loop Anchors (C), and/or Boundaries (E) and fall within expressed genes. Fisher's exact test comparing the distribution of major versus introns of interest in both expressed genes and TADs (B), Loop Anchors (D), and/or Boundaries (F).

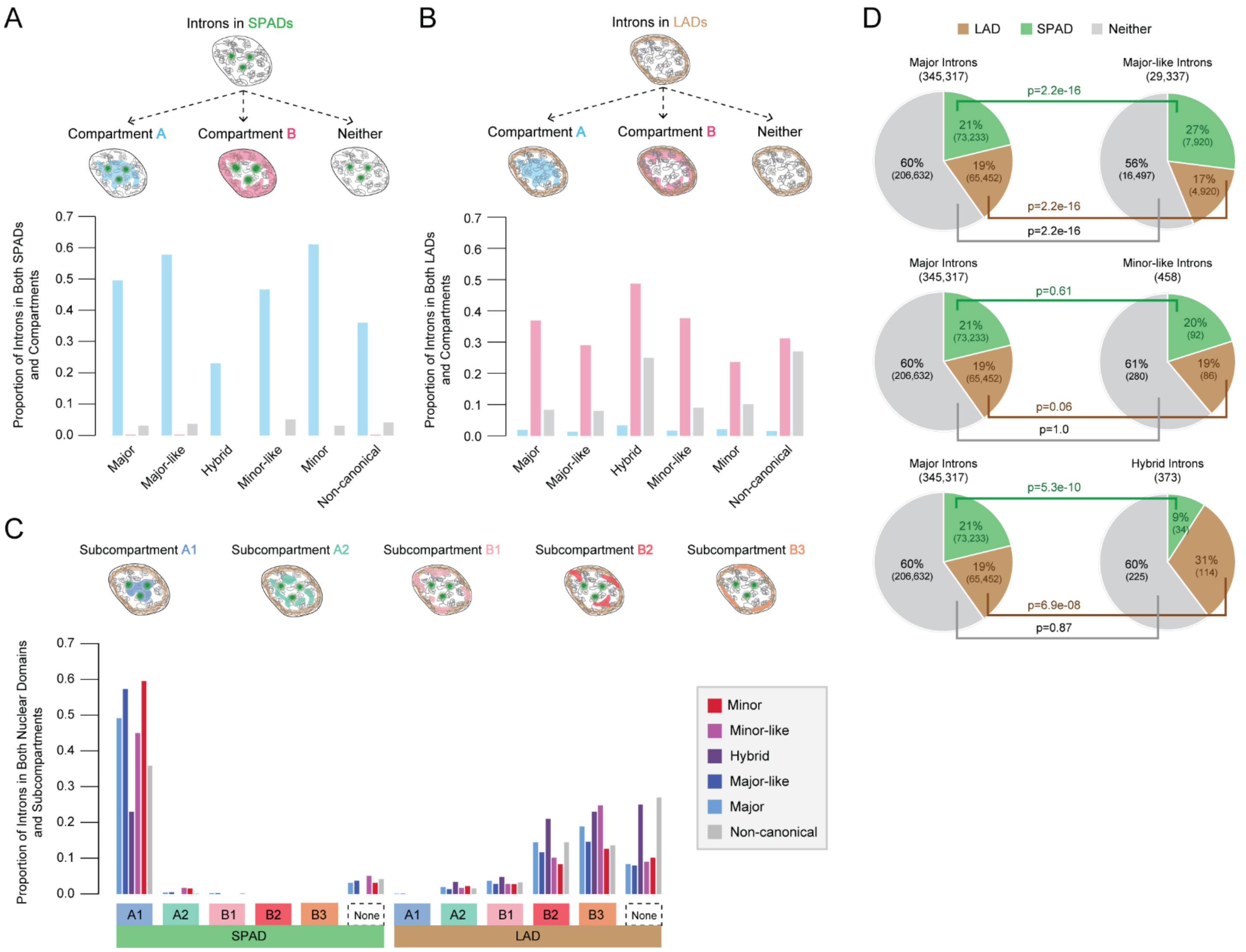


**Figure S12. Statistical analysis of remaining intron classes versus major introns in nuclear domains and intersection of introns in compartments and nuclear domains.** A. Bar graph of overlap between SPADs and compartment annotations per intron class. B. Bar graph of overlap between LADs and compartment annotations per intron class. C. Bar graph of overlap between LADs, SPADs, and subcompartment annotations per intron class. D. Fisher's exact test comparing the nuclear domain distribution of major versus major-like, minor-like, and hybrid introns.

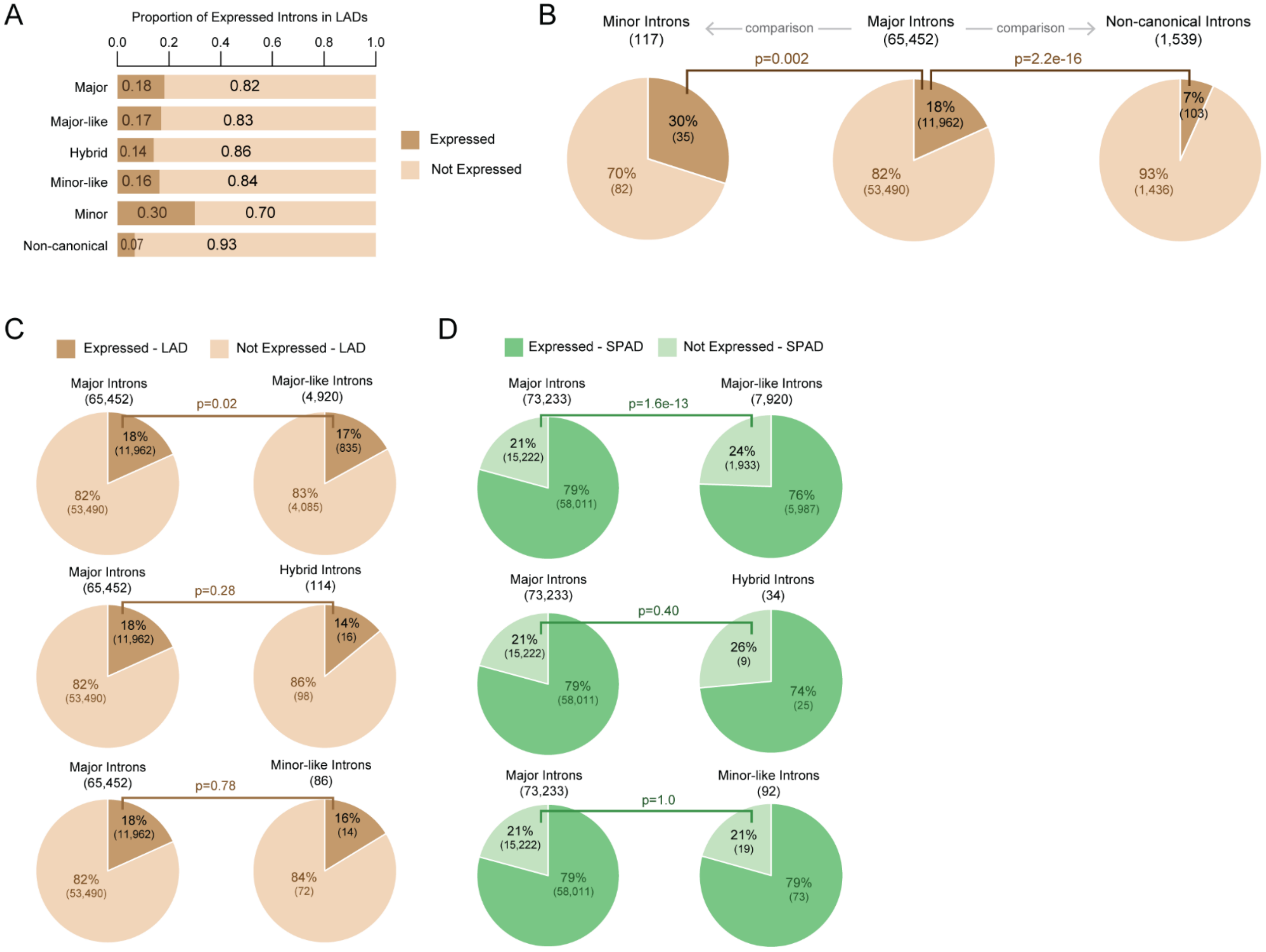


**Figure S13. Statistical analysis of remaining intron classes versus major introns in expressed genes across nuclear domains.** A. Proportion of expressed introns per class in LADs. B. Fisher's exact test comparing the distribution of major versus minor introns and major versus non-canonical introns in LADs among expressed genes. C. Fisher's exact test comparing the distribution of major versus all other intron classes in LADs and in expressed genes. D. Fisher's exact test comparing the distribution of major versus all other intron classes in SPADs and in expressed genes.

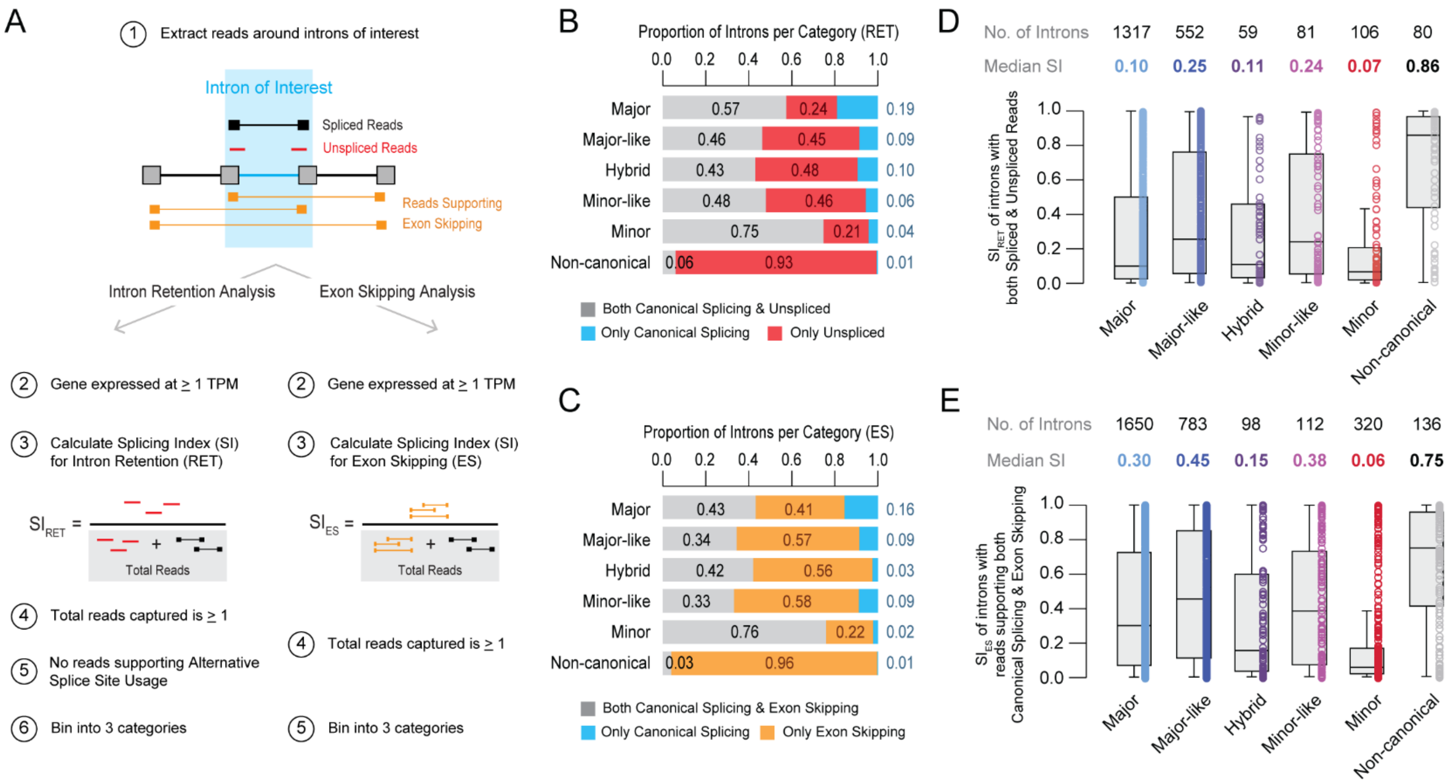


**Figure S14. Bioinformatics pipeline for intron retention and exon skipping analysis.** A. Schematic overview of splicing analysis pipeline used to quantify a splicing index (SI) for Intron Retention (RET) and Exon Skipping (ES). Only introns in genes expressed at greater than or equal to 1 TPM with at least one supporting read were leveraged for analysis. 3 Categories of Introns produced by Intron Retention and Exon Skipping analysis are described in Fig. S15. B. Proportion of introns per class with reads supporting both canonical splicing and intron retention, only canonical splicing, or only intron retention. C. Proportion of introns per class with reads supporting both canonical splicing and exon skipping, only canonical splicing, or only exon skipping. D. Distribution of $SI_{RET}$ values across intron classes for introns with reads supporting both canonical splicing and intron retention. E. Distribution of $SI_{ES}$ values across intron classes for introns with reads supporting both canonical splicing and exon skipping. Results of statistical testing to assess differences between intron retention and exon skipping between classes can be found in Table S4.

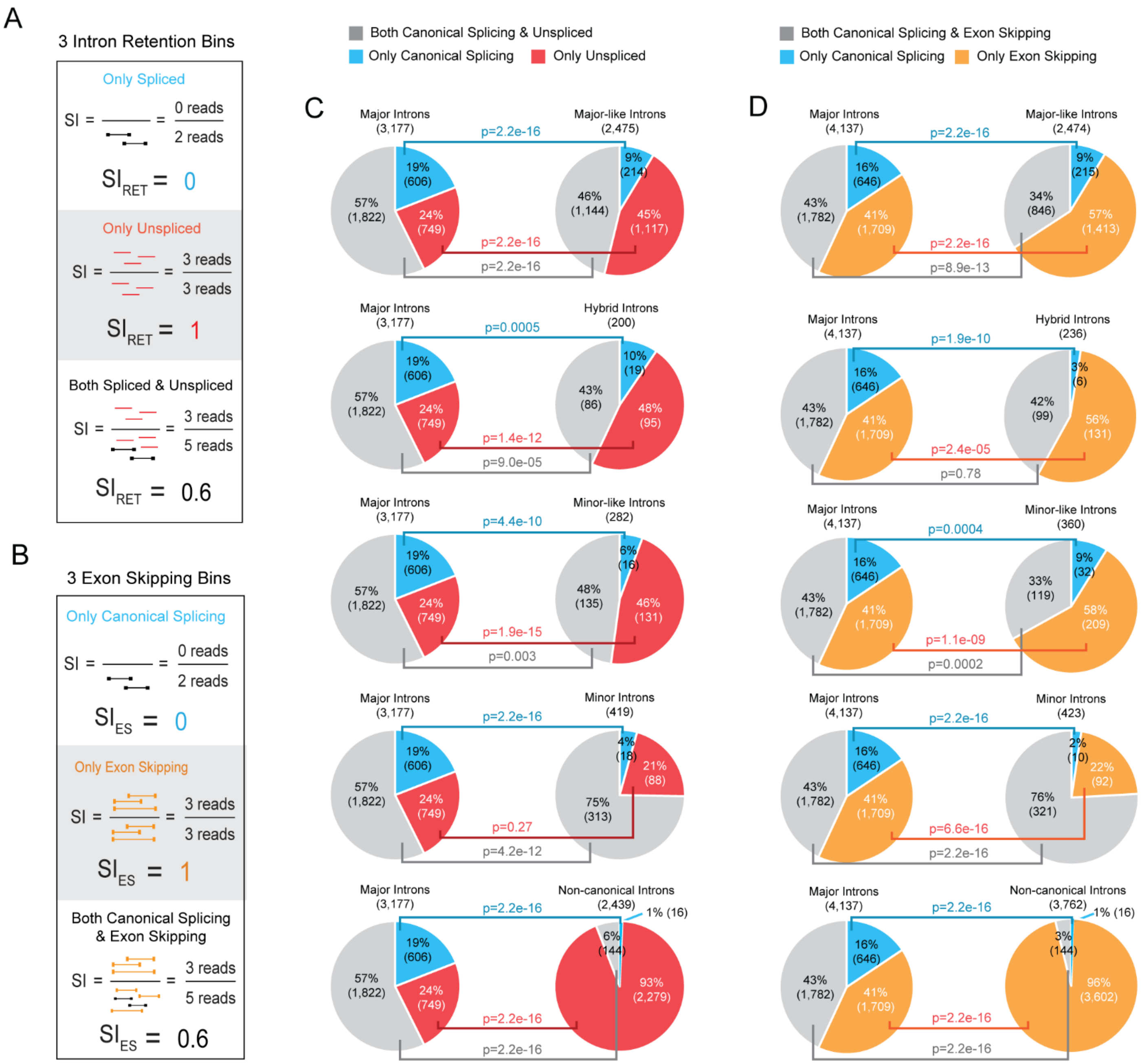


**Figure S15. Quantification of introns in exon skipping and intron retention bins.** A-B. Schematic representation of the three intron retention bins (A) and three exon skipping bins (B). C. Fisher's exact tests comparing intron retention bin distribution of major introns to each class. D. Fisher's exact tests comparing exon skipping bin distribution of major introns to each class.

**Supplementary Tables**

**Table S1.** Percentage of the coding genome occupied by major and rare intron classes.

| | Intron Occupancy in the Coding Genome | | Major Intron Occupancy | | Non-Major Intron Occupancy | | | | |
|---|---|---|---|---|---|---|---|---|---|
| | Exon % | Intron % | Major Intron % | Non-Major Intron % | Major-like % | Hybrid % | Minor-like % | Minor % | Non-canonical % |
| **All** | 3.83% | 96.17% | 93.99% | 6.01% | 5.03% | 0.07% | 0.06% | 0.11% | 0.74% |
| **1** | 4.40% | 95.60% | 94.11% | 5.89% | 4.75% | 0.02% | 0.03% | 0.15% | 0.93% |
| **2** | 3.38% | 96.62% | 94.35% | 5.65% | 4.95% | 0.18% | 0.03% | 0.07% | 0.42% |
| **3** | 2.93% | 97.07% | 93.95% | 6.05% | 4.77% | 0.05% | 0.18% | 0.31% | 0.74% |
| **4** | 2.40% | 97.60% | 95.08% | 4.92% | 4.34% | 0.02% | 0.06% | 0.07% | 0.44% |
| **5** | 2.89% | 97.11% | 95.56% | 4.44% | 3.78% | 0.04% | 0.01% | 0.07% | 0.54% |
| **6** | 3.97% | 96.03% | 94.60% | 5.40% | 4.49% | 0.08% | 0.07% | 0.10% | 0.66% |
| **7** | 4.01% | 95.99% | 93.78% | 6.22% | 5.00% | 0.07% | 0.03% | 0.09% | 1.04% |
| **8** | 3.14% | 96.86% | 94.25% | 5.75% | 5.05% | 0.08% | 0.14% | 0.04% | 0.43% |
| **9** | 3.95% | 96.05% | 94.86% | 5.14% | 3.87% | 0.02% | 0.01% | 0.11% | 1.13% |
| **10** | 3.51% | 96.49% | 95.04% | 4.96% | 4.22% | 0.04% | 0.03% | 0.06% | 0.62% |
| **11** | 6.76% | 93.24% | 94.04% | 5.96% | 5.23% | 0.00% | 0.04% | 0.04% | 0.64% |
| **12** | 4.54% | 95.46% | 93.26% | 6.74% | 5.60% | 0.13% | 0.04% | 0.06% | 0.91% |
| **13** | 3.22% | 96.78% | 93.28% | 6.72% | 5.77% | 0.05% | 0.02% | 0.09% | 0.78% |
| **14** | 3.91% | 96.09% | 93.68% | 6.32% | 5.39% | 0.01% | 0.10% | 0.15% | 0.68% |
| **15** | 4.49% | 95.51% | 92.24% | 7.76% | 6.68% | 0.08% | 0.08% | 0.09% | 0.83% |
| **16** | 4.16% | 95.84% | 94.36% | 5.64% | 4.90% | 0.06% | 0.04% | 0.11% | 0.52% |
| **17** | 6.38% | 93.62% | 92.11% | 7.89% | 6.58% | 0.06% | 0.18% | 0.07% | 1.00% |
| **18** | 3.47% | 96.53% | 93.32% | 6.68% | 6.13% | 0.01% | 0.02% | 0.10% | 0.42% |
| **19** | 9.42% | 90.58% | 92.14% | 7.86% | 6.45% | 0.05% | 0.08% | 0.12% | 1.15% |
| **20** | 5.36% | 94.64% | 93.63% | 6.37% | 4.86% | 0.10% | 0.05% | 0.14% | 1.22% |
| **21** | 3.88% | 96.12% | 94.54% | 5.46% | 4.62% | 0.31% | 0.05% | 0.04% | 0.43% |
| **22** | 7.05% | 92.95% | 91.17% | 8.83% | 7.00% | 0.14% | 0.08% | 0.23% | 1.37% |
| **X** | 2.91% | 97.09% | 93.04% | 6.96% | 5.66% | 0.13% | 0.10% | 0.16% | 0.91% |
| **Y** | 3.08% | 96.92% | 90.22% | 9.78% | 4.38% | 0.62% | 0.02% | 0.06% | 4.71% |

**Table S2.** Intron abundance per class per chromosome.

| | Major | Major-like | Hybrid | Minor-like | Minor | Non-canonical |
|---|---|---|---|---|---|---|
| **1** | 33088 | 2633 | 22 | 41 | 107 | 548 |
| **2** | 26835 | 2060 | 32 | 34 | 69 | 375 |
| **3** | 21408 | 1640 | 16 | 33 | 51 | 271 |
| **4** | 14462 | 1068 | 18 | 24 | 26 | 239 |
| **5** | 16008 | 1165 | 22 | 12 | 41 | 196 |
| **6** | 16472 | 1359 | 19 | 13 | 33 | 244 |
| **7** | 16847 | 1289 | 21 | 30 | 40 | 379 |
| **8** | 13381 | 1039 | 13 | 18 | 27 | 167 |
| **9** | 12929 | 1016 | 12 | 5 | 14 | 247 |
| **10** | 13509 | 1023 | 15 | 13 | 28 | 206 |
| **11** | 18771 | 1760 | 10 | 20 | 41 | 283 |
| **12** | 19244 | 1732 | 28 | 19 | 49 | 281 |
| **13** | 6610 | 515 | 14 | 14 | 19 | 150 |
| **14** | 11336 | 1003 | 5 | 22 | 35 | 178 |
| **15** | 13353 | 1172 | 22 | 20 | 31 | 234 |
| **16** | 15473 | 1559 | 9 | 19 | 42 | 240 |
| **17** | 19200 | 1875 | 16 | 35 | 41 | 312 |
| **18** | 6312 | 557 | 4 | 8 | 13 | 125 |
| **19** | 18420 | 2041 | 11 | 29 | 36 | 419 |
| **20** | 8189 | 683 | 12 | 14 | 21 | 104 |
| **21** | 4261 | 323 | 9 | 4 | 11 | 120 |
| **22** | 7214 | 658 | 4 | 12 | 25 | 179 |
| **X** | 10484 | 931 | 22 | 15 | 37 | 292 |
| **Y** | 1420 | 228 | 17 | 4 | 13 | 278 |

**Table S3.** Results of Fisher exact significance testing (BH adjusted p-values) on proportions of introns per class per 3D genomic feature.

| Comparison | | | Feature | | | | | |
|---|---|---|---|---|---|---|---|---|
| | | | Boundary | Loop Anchor | TAD & Loop Anchor | Loop Anchor & Boundary | TAD & Boundary | All |
| **Major** | **vs.** | **Major-like** | 0.42 | 0.33 | 0.26 | 1.0 | 0.48 | 0.56 |
| | | **Hybrid** | 1.0 | 0.71 | 0.78 | 0.41 | 1.0 | 1.0 |
| | | **Minor-like** | 0.21 | 1.0 | 1.0 | 1.0 | 0.63 | 1.0 |
| | | **Minor** | 0.27 | 0.73 | 0.44 | 0.14 | 0.47 | 0.62 |
| | | **Non-canonical** | 0.11 | **1.7e-08** | **5.9e-05** | **0.0004** | 0.14 | **0.005** |

**Table S4.** Statistical analysis results of Kruskal-Wallis test to assess differences in exon skipping and intron retention between intron classes, regardless of 3D location.

| Comparison | | | Adj. p-value for Intron Retention | Adj. p-value for Exon Skipping |
|---|---|---|---|---|
| Hybrid | vs. | Minor | 8.65E-02 | 7.80E-05 |
| Hybrid | | Major | 5.54E-01 | 1.08E-01 |
| Minor | | Major | **4.39E-02** | **4.09E-25** |
| Hybrid | | Major-like | 1.28E-02 | 3.95E-04 |
| Minor | | Major-like | 6.58E-08 | 7.27E-37 |
| Major | | Major-like | 2.55E-08 | 1.78E-06 |
| Hybrid | | Minor-like | 8.05E-02 | 8.77E-02 |
| Minor | | Minor-like | 7.66E-05 | 1.89E-10 |
| Major | | Minor-like | 1.12E-03 | 4.37E-01 |
| Major-like | | Minor-like | 5.56E-01 | 1.85E-01 |
| Hybrid | | Non-canonical | 2.77E-08 | 4.89E-10 |
| Minor | | Non-canonical | 2.67E-17 | 1.18E-36 |
| Major | | Non-canonical | 2.09E-19 | 2.45E-13 |
| Major-like | | Non-canonical | 2.88E-05 | 1.82E-06 |
| Minor-like | | Non-canonical | 3.26E-05 | 5.63E-06 |

**Supplementary Data**

**Dataset S1.** Intron abundance per 250 kb bin. (csv format). This table reports the coordinates and intron content of every 250 kb bin created across the human genome for the quantification of intron density. Intron numbers per intron class and bin are reported in addition to the corresponding parent genes.

**Dataset S2.** Hi-C and other 3D dataset annotation information per cell line. (csv format). This table reports the dataset availability for 3D features in K562 cells and cell line data for SPAD and LAD annotations used in the present study. Basic data annotations such as the number of regions, median region size (bp), coverage (bp), and portion of the genome covered (as a percentage) are also reported.

**Dataset S3.** K562 loop anchor annotations. (csv format). Provides the genomic coordinates and supporting information of loop anchor annotations produced from K562 Hi-C data leveraged in the current manuscript.

**Dataset S4.** K562 TAD annotations. (csv format). Provides the genomic coordinates and supporting information of TAD annotations produced from K562 Hi-C data leveraged in the current manuscript.

**Dataset S5.** Intersects of introns with 3D genomic features in K562, HCT116, HFFc6, and H1 cells requiring 50% intron overlap with each feature. (csv format). Provides intersection data for 3D features from four cell lines with introns from six classes when a minimum of 50% overlap is required. 1 indicates a positive intersection while 0 indicates no overlap.

**Dataset S6.** Intersects of introns with 3D genomic features in K562 cells requiring 1 bp or 90% intron overlap with each feature. (csv format). Provides intersection data for 3D features with introns from six classes when either a minimum of 1 bp or 90% overlap is required. 1 indicates a positive intersection while 0 indicates no overlap.

**Dataset S7.** List of genes in the total and core essentialomes. (csv format). Lists all genes in the total and core essentialomes as previously annotated by Meyers et al. and interrogated in Baumgartner et al. (14, 15).

**Dataset S8.** RNA-seq dataset information per cell line. (csv format). This table reports the dataset availability for transcriptome data in K562, H1, HCT116, and HFFc6 cells. Information includes sample names, study first author, PMID, GEO Accession number, and associated SRA run identifiers (SRR numbers).

**Dataset S9.** List of expressed genes per cell line. (csv format). This table lists all genes identified as expressed (greater than or equal to 1 TPM) in K562, H1, HCT116, and/or HFFc6 cells based on RNA-seq analysis.

**Dataset S10.** Subcompartment expression by intron class. (csv format). Provides the proportion values for introns in expressed and not-expressed genes per subcompartment per intron class. Also included are differences in proportions of introns in expressed and not-expressed genes per intron class, corresponding to Fig. S10F.

**Dataset S11.** Summary of statistical comparison of intron retention across intron classes and cell lines. (csv format). This table reports adjusted p-values for an all to all comparison of intron retention across intron classes and cell lines. Corresponding significance symbols are displayed in the far right of the table.

**Dataset S12.** Summary of statistical comparison of exon skipping across intron classes and cell lines. (csv format). This table reports adjusted p-values for an all to all comparison of exon skipping across intron classes and cell lines. Corresponding significance symbols are displayed in the far right of the table.

**Dataset S13.** GO terms with parent genes of introns with increased splicing efficiency in K562 and HCT116 relative to HFFc6 cells. (csv format).
